# Galectin-1 Modulates Cell Adhesions, Caveolae, and Vascular Permeability in Kidney Endothelial Cells – Insights from Proteomics, Phosphoproteomics, and Functional Studies

**DOI:** 10.64898/2026.03.03.709385

**Authors:** Alex Boshart, Sofia Farkona, Shravanthi Rajasekar, Chiara Pastrello, Maya Allen, Stefan Petrovic, Kieran Manion, Slaghaniya Neupane, Sergi Clotet-Freixas, Hong Sang Choi, Anne-Marie Bulboaca, Rohan John, Allen Duong, Stephen Juvet, Milica Radisic, Juan Manuel Pérez Sáez, Gabriel A. Rabinovich, Sara Deir, Boyang Zhang, Igor Jurisica, Ana Konvalinka

**Affiliations:** Toronto General Hospital Research Institute, University Health Network, Toronto, ON, Canada; Ajmera Transplant Centre, University Health Network, Toronto, ON, Canada; Institute of Medical Science, University of Toronto, Toronto, ON, Canada; Osteoarthritis Research Program, Division of Orthopedic Surgery, Schroeder Arthritis Institute, University Health Network, Toronto, ON, Canada; Data Science Discovery Centre for Chronic Diseases, Krembil Research Institute, University Health Network, Toronto, ON, Canada; Department of Laboratory Medicine and Pathobiology, University of Toronto, Toronto, ON M5S 1A8, Canada; Division of Nephrology, McMaster University and St. Joseph’s Healthcare, Hamilton, Ontario, Canada; Department of Internal Medicine, Chonnam National University Medical School and Hospital, Gwangju, Republic of Korea; Institute of Biomedical Engineering, University of Toronto, Toronto, ON, Canada; Department of Chemical Engineering and Applied Chemistry, University of Toronto, Toronto, ON, Canada; Laboratory of Glycomedicine, Instituto de Biología y Medicina Experimental (IBYME), Consejo Nacional de Investigaciones Científicas y Técnicas (CONICET), 1428 Ciudad de Buenos Aires, Argentina; Facultad de Ciencias Exactas y Naturales, Universidad de Buenos Aires, 1428 Ciudad de Buenos Aires, Argentina; Laboratory of Glycoimmunology, Caixa Research Institute, 08022 Barcelona, Spain; Department of Chemical Engineering, McMaster University, Hamilton, Ontario, L8S 4L8 Canada; School of Biomedical Engineering, McMaster University, Hamilton, Ontario, L8S 4L8 Canada; Departments of Medical Biophysics and Computer Science, and Faculty of Dentistry, University of Toronto, Toronto, ON, Canada; Institute of Neuroimmunology, Slovak Academy of Sciences, Bratislava 845 10, Slovakia; Department of Medicine, Division of Nephrology, University Health Network, Toronto, ON, Canada

**Keywords:** Primary glomerular microvascular endothelial cells, Galectin-1, Interferon-gamma, Proteomics, Phosphoproteomics, siRNA knockdown, Antibody mediated rejection

## Abstract

Microvascular inflammation and endothelial injury, triggered by interferon-gamma (IFNγ), are hallmarks of antibody-mediated rejection (ABMR), the leading cause of premature kidney allograft loss. Glomerular extracellular matrix (ECM) remodeling and endothelial caveolae formation are important aspects of chronic ABMR. We found galectin-1, an immunomodulatory protein that interacts with the ECM, to be increased in the glomeruli of patients with ABMR, while its gene (*LGALS1*) expression was decreased by IFNγ stimulation in glomerular endothelial cells. Mechanisms underlying endothelial dysfunction in ABMR, its links to ECM remodeling, and the role of immunomodulatory proteins such as galectin-1 remain incompletely understood. Here we studied the effects of galectin-1 modulation in glomerular microvascular endothelial cells (GMECs) *in vitro*. We demonstrated that galectin-1 was mainly expressed by glomerular endothelial cells in ABMR kidneys. To model key aspects of endothelial injury in ABMR, we knocked down *LGALS1* in GMECs, followed by stimulation with IFNγ and performed label-free quantitative proteomic and phosphoproteomic profiling of GMECs. Proteomic analysis identified 5446 proteins (FDR<0.01), of which 236, 827, and 267 were differentially expressed in response to *LGALS1* knockdown, IFNγ treatment, and their interaction, respectively (FDR<0.05). Both *LGALS1* knockdown and the interaction between treatments significantly altered expression of adhesion proteins (FDR<0.01), particularly integrin subunit β5, which was validated. Phosphoproteomic profiling identified 2727 phosphopeptides (FDR<0.01), with 28 that were differentially expressed across *LGALS1* knockdown, IFNγ treatment, and their interaction (P<0.01). Phosphorylation of CAVN1 and co-localization with its partner CAV1, critical for caveolar formation, were decreased in GMECs upon *LGALS1* knockdown, IFNγ stimulation, or both. In a microfluidic model of the glomerular microvasculature, addition of recombinant galectin-1 increased both endothelial permeability and secretion of proinflammatory cytokines, in *LGALS1*-silenced GMECs. Thus, endothelial signaling pathways regulated by inflammatory cues and galectin-1 contribute to endothelial injury and caveolae formation, highlighting galectin-1 as a potential therapeutic target in ABMR.

**Synopsis:** Galectin-1 is expressed by kidney glomerular endothelium. This study reveals that modifying galectin-1 in endothelial cells, in the presence of IFNγ perturbs cytoskeletal, adhesion and caveolar proteins resulting in altered endothelial permeability.

- *LGALS1* knockdown increased ECM proteins and decreased interferon-induced proteins.
- *LGALS1* knockdown and IFNγ treatment perturbed cell adhesion proteins such as ITGB5.
- CAVN1 phosphorylation and colocalization with CAV1 decreased upon *LGALS1* knockdown.
- Extracellular galectin-1 increased microvascular permeability in response to IFNγ.

## INTRODUCTION

Kidney disease is a global health problem, affecting over 14% of the general population worldwide (Mark *et al*, 2025). Kidney transplantation is the best treatment for patients with end-stage kidney disease (Pascual *et al*, 2002; Hariharan *et al*, 2021; Wolfe *et al*, 1999), which carries mortality comparable to terminal cancer (Tonelli *et al*, 2022). However, over 50% of kidney transplant recipients lose their grafts by 20 years post-transplant. The main cause of premature kidney allograft loss is antibody-mediated rejection (ABMR) (Gaston *et al*, 2010; Sellarés *et al*, 2012). ABMR is driven by donor-specific antibodies (DSAs) that target human leukocyte antigen (HLA) expressed on the allograft endothelium (Patel & Terasaki, 1969; Hirohashi *et al*, 2012; Konvalinka & Tinckam, 2015). These interactions can initiate a cascade of complement activation, endothelial activation, and leukocyte recruitment, ultimately leading to microvascular inflammation, tissue injury, and graft dysfunction (Garg *et al*, 2017; Blume *et al*, 2012). Glomerular extracellular matrix (ECM) remodelling is a cardinal feature of chronic ABMR, and many native glomerular diseases, but the potential contributions of the microvascular endothelium to this ECM remodelling, and the drivers of endothelial phenotypic change remain incompletely understood (Novotná *et al*, 2021; Rodriguez-Ramirez *et al*, 2022).

Pro-inflammatory cytokines, such as interferon-gamma (IFNγ), are secreted by infiltrating immune cells in the allograft microenvironment (O’Neill & Hidalgo, 2021; Zhang *et al*, 2024; Boshart *et al*, 2024). IFNγ upregulates endothelial class II HLA expression which, in the presence of DSA, can promote antibody induced injury through Fcγ receptors or complement activation (Collins *et al*, 1984; Wedgwood *et al*, 1988; Xie *et al*, 2019). In our previous proteomic analysis of ABMR biopsies, we found that galectin-1 (encoded by the *LGALS1* gene), a small, evolutionarily conserved, carbohydrate-binding protein, was increased in the glomeruli of ABMR patients, while ECM proteins interacting with galectin-1 were decreased, compared to biopsies with different types of graft injury (Clotet-Freixas *et al*, 2020). Further, we demonstrated that universal anti-HLA class I antibodies and IFNγ altered transcriptional expression and protein secretion of galectin-1 in kidney glomerular microvascular endothelial cells (GMECs) (Clotet-Freixas *et al*, 2020). Remarkably, galectin-1 is an immunomodulatory protein that can impair the functionality of cytotoxic T-cells through interaction with various T-cell surface glycoproteins including CD45 and several ECM glycoproteins such as laminin and fibronectin (Perillo *et al*, 1997; Stillman *et al*, 2006; Ozeki *et al*, 1995; Qun & Cummings, 1993). Preclinical models have shown the potential of this lectin to prolong allograft survival in kidney and skin transplantation (Xu *et al*, 2010; Moreau *et al*, 2012; Alhabbab *et al*, 2018). Finally, both the addition of galectin-1 and its inhibition modulate immune cell adhesion and extravasation through the endothelium and ECM (RABINOVICH *et al*, 1999; Norling *et al*, 2008; Nambiar *et al*, 2019). We propose that modulation of galectin-1 in human GMECs, particularly when exposed to IFNγ, would alter integrity and functionality of the endothelial barrier.

In this study, we performed label-free quantitative proteomic and phosphoproteomic profiling of primary human GMECs to define the effect of *LGALS1* knockdown in the context of IFNγ stimulation, thus modelling functional patterns associated with endothelial cell activation, characteristic of ABMR. Our analyses revealed that both IFNγ stimulation and *LGALS1* knockdown induced coordinated changes in proteins and phosphorylation sites associated with focal adhesion dynamics and cytoskeletal remodeling, which are critical for endothelial barrier function, immune cell interaction, and mechano-transduction. We also assessed the functional effects of *LGALS1* knockdown and addition of recombinant galectin-1 on GMEC adhesion and permeability. These findings provide new insights into the regulation of endothelial cell structure and signaling in response to inflammatory cytokines and galectin-1, offering potential therapeutic targets to mitigate microvascular injury in kidney transplantation, and more broadly, in glomerular kidney disease.

## MATERIALS AND METHODS

### Experimental Design and Statistical Rationale

Four experimental conditions were analyzed to assess the proteomic and phosphoproteomic changes induced by *LGALS1* knockdown (siRNA) and IFNγ treatment: **(1)** *LGALS1* siRNA + IFNγ, **(2)** *LGALS1* siRNA + vehicle, **(3)** non-targeting control + IFNγ, and **(4)** non-targeting control + vehicle. Experiments were performed on separate days using new lots of thawed GMECs, to create an experimental unit of 3 (N = 3) for each condition, resulting in a total of 12 samples per experiment for both proteomic and phosphoproteomic analyses. This number of biological replicates was determined to be adequate to identify proteins and phosphopeptides that are differentially expressed in at least two out of three biological experiments, in prior similar experiments (Clotet-Freixas *et al*, 2020; Clotet *et al*, 2017; Konvalinka *et al*, 2013; Clotet-Freixas *et al*, 2024; Rajasekar *et al*, 2022; Didangelos *et al*, 2010; Okanishi *et al*, 2022).

The duration of IFNγ treatments selected for the phosphoproteomics or proteomics experiments were 30 minutes or 24 hours, respectively. At these time points, IFNγ treatment resulted in maximal STAT1 phosphorylation (30 minutes) (**Supplemental Figure S1**) and increased expression of HLA class II (24 hours) after IFNγ signalling (**Supplemental Figure S2**) and is consistent with prior literature (Clotet-Freixas *et al*, 2020; Blanchette *et al*, 2003; Marchetti *et al*, 2006; Yang *et al*, 2005; Lee *et al*, 2021; Mutthi *et al*, 2018).

### Cell culture of GMECs

Commercial primary human GMECs were purchased from Cell Systems and cultured in Endothelial Cell Growth Media MV (C-22120; Promocell), supplemented with the ready-to-use kit containing 10% (v/v) dialyzed fetal calf serum (FCS), 10 ng/mL epidermal growth factor (EGF), 1 µg/mL hydrocortisone, and 90 µg/mL heparin. Penicillin (50 U/mL) and streptomycin (50 µg/mL) were added to the medium. Experiments were conducted using cells between passages 6 and 8.

### Electroporation for siRNA *LGALS1* knockdown and IFNγ treatment

GMECs at passage 6 were electroporated at 1 million cells per cuvette using the P5 Primary Cell 4D-Nucleofector X Kit (Lonza) and program CA-167 on a 4D-Nucleofector device (Lonza). Program settings and cell viability were assessed after electroporation using the green fluorescent protein (GFP) vector included in the Nucleofector X Kit and flow cytometry. Next, cells were transfected with 100 nM siRNA SmartPool targeting *LGALS1* (Dharmacon) or non-targeting control siRNA (Dharmacon). Knockdown efficiency was confirmed on the mRNA level by RT-qPCR and on the protein level by LC-MS/MS or ELISA.

To assess IFNγ-induced cell signalling changes in the presence or absence of *LGALS1*, 6-days after electroporation, when GMECs achieved 100% confluence, cells were serum-starved for 18 hours. They were then treated with 50 U/mL of IFNγ (SRP3058; Sigma) or vehicle containing 0.1% BSA in PBS for 30 minutes and collected as cell pellets, snap frozen, and stored at −80°C until further phosphoproteomic analyses.

To assess protein-level changes due to IFNγ in the presence or absence of *LGALS1*, 5-days after electroporation, when GMECs achieved 100% confluence, cells were serum-starved for 18 hours. They were then treated with 50 U/mL of IFNγ (SRP3058; Sigma) or vehicle containing 0.1% BSA in PBS for 24 hours and collected as cell pellets, snap frozen, and stored at −80°C until further proteomic analyses.

### Proteomics

Lysis and digestion: Cell pellets were processed for LC-MS/MS as previously described (Clotet *et al*, 2017). Briefly, cell pellets were resuspended in 200µL of 0.1% w/v acid-labile detergent RapiGest SF (Waters) in 50 mM ammonium bicarbonate. Samples were vortexed and sonicated with a tip probe (three 30 second cycles at 80% output power) (Fisher Scientific) followed by centrifugation for 20 min at 15,000g at 4°C. Lysates were transferred to clean microcentrifuge tubes and total protein concentration was measured using the Micro BCA Protein Assay Kit (Thermo Scientific). Next, 100µg of total protein from each sample/lysate were reduced with 10 mM dithiothreitol at 60°C for 15 min and alkylated with 20mM of iodoacetamide at room temperature for 15 min, protected from light. Samples were then mixed with trypsin (1:50 ratio) and incubated overnight (18 hours) at 37°C, to digest the proteins into peptides. The next day, 1% trifluoroacetic acid was added to each sample to cleave the RapiGest SF detergent and samples were vortexed and centrifuged for 15min at 15,000g and 4⁰C. Supernatants containing the peptides were then transferred to clean microcentrifuge tubes.

LC-MS/MS analysis: Peptides were desalted and concentrated with OMIX C18 tips (Agilent) and loaded in technical duplicates onto a 3cm long C18 precolumn (75μm ID packed with Agilent Pursuit 5μm beads) and a C18 EASY-Spray™ 2μm, 500mm × 75μm capillary column (Thermo Fisher Scientific) using EASY-nLC 1200 pump (Thermo Fisher Scientific, San Jose, CA) at 20μL/min. Mobile phases included 0.1% formic acid in water (buffer A) and 0.1% formic acid in 95% acetonitrile (buffer B). Peptides were separated on a 135 min gradient elution at 250 nL/min. A four-step gradient was used: 3% to 30% of buffer B for 110 min, 30% to 65% for 10 min, 65% to 100% for 5 min and 100% for 10 min at 300 nL/min. The liquid chromatography setup was coupled online to a Q-Exactive™ HF-X Hybrid Quadrupole-Orbitrap Mass Spectrometer (Thermo Fisher Scientific) with a nanoelectrospray ionization source. The peptides were analyzed in a data-dependent mode with one survey scan with automatic gain control (AGC) of 3 x 10^6^ ions (400-1500 m/z, R=60,000) followed by top 10 MS/MS scans with high-energy collisional dissociation-based fragmentation (AGC 1x10^5^ ions, maximum filling time 22ms, isolation window 1.6 m/z and normalized collision energy 27) detected in the Orbitrap (R=15,000).

Proteomics data analysis: The acquired raw data files were processed using MaxQuant software (Ver.1.6.0.16) with Andromeda search engine (Cox & Mann, 2008). The data were searched using a target-decoy approach with a reverse database against UniProt Human reference proteome fasta file (downloaded from UniprotKB on Nov 19, 2024) with an FDR<1% at the level of proteins, peptides, and modifications. The default settings were used and match between runs was enabled. MaxQuant ‘ProteinGroups’ file from the proteome run was imported into Perseus Ver.2.0.11 (Tyanova *et al*, 2016). Results were filtered using the reverse hits function in Perseus and known contaminants were removed. Technical replicates were assessed for missing values, and the replicate with the higher number of missing values was removed. Proteins were retained if they were found in >30% of the samples. Multiple different cutoffs were tested, and the number of total proteins diminished by only ∼200 proteins at this cutoff. Missing values were then imputed (**Supplemental Figure S3)**. Batch correction was performed using the Perseus plugin ComBat in R Ver.4.4.2. The statistical analysis was performed using a 2-way ANOVA and Tukey’s Post Hoc Test. Proteins were considered significant with an adj.p-value<0.05. Unsupervised hierarchical clustering of significant proteins was performed in R (Ver.4.2.3). Significant proteins were analyzed using Gene Ontology Enrichment and Pathway Enrichment Analyses (FDR:BH<0.01) (Pastrello *et al*, 2024; Ashburner *et al*, 2000; Thomas *et al*, 2022; The Gene Ontology Consortium, 2026). The mass spectrometry proteomics data have been deposited to the ProteomeXchange Consortium via the PRIDE (Perez-Riverol *et al*, 2024) partner repository with the dataset identifier PXD074919.

### Phosphoproteomics

Lysis and digestion: GMECs were processed as previously described (Humphrey *et al*, 2018). Cells were washed with TBS, resuspended in lysis buffer (4% w/v SDC, 100mM Tris pH 8) and heated at 95°C for 5 min. Following sonication with a tip probe (three 30-second cycles of 1 second on and 1 second off at 80% output power) (Fisher Scientific) and total protein concentration measurement (BCA), 200µg from each sample/lysate were treated with reduction/alkylation buffer (100 mM TCEP and 400mM 2-chloroacetamide, pH 7-8). Samples were then mixed with LysC and trypsin (1:100 ratio, wt/wt) and incubated overnight (18h) at 37°C with shaking/agitation (1,500 r.p.m) (Labnet).

Phosphopeptide enrichment: Digested peptides were mixed with 400 µL isopropanol (1,500 r.p.m., 30 seconds), followed by 100 µL of Enrichment Protocol (EP) buffer (48% [v/v] TFA, 8mM KH₂PO₄), and mixed again. Any precipitation was cleared by centrifugation at 2000g for 15min at room temperature, and the resulting supernatants were transferred to clean microcentrifuge tubes. For the phosphopeptide enrichment step, TiO_2_ beads (prepared in 6% TFA buffer, 80% ACN) were added into each sample at a 1:12 (wt/wt) protein-to-beads ratio and incubated at 40°C with agitation (2,000 r.p.m.) for 5 min. TiO_2_-bound phosphopeptides were pelleted by centrifugation (2,000g for 1 min at RT) and transferred to clear tubes while the non-phosphopeptide supernatant was stored for the whole proteome analysis. TiO_2_ beads were washed 5 times with a buffer consisting of 5% (v/v) TFA and 60% (v/v) isopropanol and transferred on top of an AttractSPE C8 tip (Affinisep) and centrifuged (1,500g for ∼8 min at room temperature) until dryness. The phosphopeptides were obtained with elution buffer (20% NH_4_OH, 40% ACN) and concentrated in a CentriVap Speed-Vac (Labconco) for approximately 20 minutes at 45⁰C. Phosphopeptides were acidified with 100μL 1% of TFA and were loaded onto an equilibrated AttractSPE SDB RPS tip (Affinisep) for desalting and further clean up. Peptides were then washed with 1% TFA in isopropanol and further washed with 0.2% v/v TFA and 5% v/v ACN. Finally, they were eluted with 60μL elution buffer (60% ACN, 1.25% NH4OH). After drying under vacuum, phosphopeptides were resuspended in MS loading buffer (2% ACN, 0.3% TFA). Total proteome containing supernatant was dried in a SpeedVac and resuspended in 2% ACN, 0.3% TFA.

LC-MS/MS analysis: Phosphorylated and non-phosphorylated peptides were desalted with OMIX C18 tips (Agilent) and loaded onto a 3cm long C18 precolumn and liquid chromatography setup was coupled online to a Q-Exactive™ HF-X Hybrid Quadrupole-Orbitrap Mass Spectrometer (Thermo Fisher Scientific) with a nanoelectrospray ionization source, as described in the Proteome Methods above.

Phosphoproteomics data analysis: The acquired raw data files from the TiO_2_-enriched runs and the non-enriched runs were processed separately using MaxQuant software (Ver.1.6.0.16) with Andromeda search engine (Cox & Mann, 2008). The data were searched using a target-decoy approach with a reverse database against UniProt Human reference proteome fasta file (downloaded from UniprotKB on Feb 11, 2025) with an FDR<1% for proteins, peptides, and modifications. The default settings were used, except for variable modifications, where oxidized methionine (M), acetylation (protein N-term) and phospho (STY) were selected for the enriched runs whereas only oxidized methionine (M) and acetylation (protein N-term) were selected for the non-enriched runs. Carbamidomethyl (C) was chosen as a fixed modification and match-between-runs was enabled. The rest of the MaxQuant settings were default. MaxQuant evidence tables from phosphopeptide-enriched runs and the corresponding non-enriched proteome runs were imported into the MSstatsPTM R package (Ver.2.8.1) for label-free analysis (Kohler *et al*, 2023). Conversion to MSstats-compatible inputs was performed with the MaxQtoMSstatsPTMFormat function, which generated phosphorylation (site-level) and protein (background) tables containing position information for each phosphorylation site, relative to the protein sequence. Peptide intensities were then processed with the dataSummarizationPTM function which performed imputation of missing values, normalization across runs using the equalize medians method, and robust summarization of peptide abundances with Tukey’s median polish (TMP). Finally, the groupComparisonPTM function was used to fit separate statistical models for phosphorylation and protein summaries and to perform differential phosphorylation analysis. Estimates of differential abundance were then adjusted for abundances in the corresponding peptide levels, resulting in adjusted fold-change estimates. Phosphopeptides that could not be adjusted to their unmodified peptide counterpart or resulted in an adjusted fold-change estimate of infinity were removed. Phosphopeptides were considered significant with an FDR<0.2 after comparison to their unmodified peptide using the Benjamini Hochberg method. The mass spectrometry phosphoproteomics data have been deposited to the ProteomeXchange Consortium via the PRIDE(Perez-Riverol *et al*, 2024) partner repository with the dataset identifier PXD074969.

### Peripheral blood mononuclear cells (PBMCs) processing

Peripheral blood mononuclear cells (PBMCs) were isolated from whole blood collected from healthy volunteers. Blood was processed using EasySep (Stem Cell Technologies) with SepMate tubes (Stem Cell Technologies) following manufacturer’s protocol. Isolated cells were frozen at −80°C then stored in LN2 until use. For experiments, cells were thawed and counted in RPMI 1640 medium (Gibco, Cat#11875093) supplemented with 10% FBS (Gibco, Cat#16140071).

### Western-blotting

GMEC pellets were solubilized in 10x lysis buffer (Cell Signaling, #9803) by mechanical homogenization. Protein concentration was measured using the Micro BCA Protein Assay Kit (Thermo Scientific). For Western blot analysis of protein expression, 15-18µg total protein was loaded onto 10% acrylamide gels (Bio-Rad), separated by sodium dodecyl sulfate-polyacrylamide gel electrophoresis (SDS-PAGE), and transferred to polyvinylidene difluoride membranes (Millipore). Membranes were blocked with 5% milk and incubated overnight at 4°C with rabbit monoclonal anti-HLA-DPB1 antibody (1:2000; ab157210; Abcam) or rabbit polyclonal anti-β5 integrin (ITGB5) (1:3000; 28543-1-AP; Proteintech) followed by HRP-conjugated anti-rabbit secondary antibody (1:4000; A0545; Sigma). Membranes were stripped and incubated with mouse polyclonal anti-vinculin antibody (1:2000; MAB6896; R&D Systems) or mouse monoclonal anti-GAPDH (1:2000; CB1001; Sigma) and corresponding HRP-conjugated anti-mouse secondary antibody (1:4000; P0447; Dako) as a loading control. Chemiluminescent reaction was induced using the ECL kit (Bio-Rad, Cat #170-5060) and detection was performed using a Bio-Rad Gel Doc XR+ Imaging System.

### Immunofluorescence

GMECs were grown to confluence on coverslips placed in 6-well plates. Coverslips were washed once with 1 mL warm phosphate-buffered saline (PBS) and fixed with 1.5 mL ice-cold 4% formaldehyde (Thermo Scientific) per well for 20 minutes. After three washes with room temperature PBS, cells were permeabilized with 0.1% Triton X-100 and blocked with 7.5% normal goat serum for 1 hour. Samples were incubated overnight at 4°C with rabbit polyclonal anti-galectin-1 antibody (1:100; GTX101566; GeneTex), rabbit polyclonal anti-CAVN1 antibody (1:1000; 18892-1-AP; Proteintech), or mouse monoclonal anti-CAV1 antibody (1:1000; 66067-1-IG; Proteintech). After washing, cells were incubated with Alexa Fluor 647-conjugated anti-rabbit secondary antibody (1:200; A-21245, Invitrogen), CF595-conjugated anti-rabbit secondary antibody (1:500; SAB4600107; Sigma), or FITC-conjugated anti-mouse secondary antibody (1:500; F0257; Sigma) for 1 hour at room temperature. Cytoskeleton was stained using ActinRed 555 (Invitrogen), and nuclei were stained with 4′,6-diamidino-2-phenylindole (DAPI, Thermo Scientific). Coverslips were mounted onto slides and images were acquired using a Leica TCS SP8 confocal microscope (Leica Microsystems).

### Quantitative RT-PCR

GMECs were harvested at confluence following treatment and stored lysates in RLT buffer (Qiagen) at −80°C until RNA extraction using the RNeasy Mini Kit (Qiagen). RNA concentration was determined using a Nanodrop spectrophotometer (Thermo Scientific). Between 300 and 1800 ng mRNA was reverse-transcribed to cDNA using the High-Capacity cDNA Reverse Transcription Kit (Applied Biosystems). Transcript levels of *LGALS1, CDH5, PECAM1, VWF, ACTA2, IL-6,* and *ICAM1* were quantified by real-time PCR using Power SYBR Green PCR Master Mix (Applied Biosystems) on a LightCycler 480 Real-Time PCR System (Roche). Transcript levels were normalized to the housekeeping gene β-actin (*ACTB*). Primer sequences are provided in **Supplemental Table S1**.

### Flow cytometry

Flow cytometry was used to assess intracellular galectin-1 expression in both human PBMCs and GMECs. Both PBMCs and GMECs were thawed, washed with phosphate-buffered saline (PBS), and resuspended in FACS buffer (PBS containing 2% FCS and 2 mM ethylenediaminetetraacetic acid (EDTA)). Cell viability was assessed using LIVE/DEAD Fixable Near-IR Dead Cell Stain (LIVE/DEAD 700; Thermo Fisher) according to the manufacturer’s instructions. After staining, cells were washed twice with FACS buffer and fixed and permeabilized with BD Cytofix/Cytoperm Kit (BD Biosciences) according to the kit instructions. Cells were then stained intracellularly with FITC-conjugated anti-human galectin-1 antibody (clone GAL1/1831; Novus Biologicals) for 30 minutes at 4°C. Fluorescence minus one (FMO) control was used for galectin-1 in both PBMC and GMEC analyses to establish gating thresholds. Single-color compensation was performed using UltraComp eBeads (Thermo Fisher) and Amine Reactive Compensation Bead Kit (Thermo Fisher) for each fluorophore, including viability dye. Samples were acquired on a BD LSRFortessa flow cytometer (BD Biosciences), and data were analyzed using FlowJo software (v10, BD Biosciences). Galectin-1 expression was quantified as median fluorescence intensity (MFI) of galectin-1 positive cells within each gated cell type.

### ELISA

To evaluate galectin-1 secretion into the GMEC cell media, GMEC supernatants were collected post-electroporation and analyzed in duplicate using the Human galectin-1 ELISA Kit (KA5065; Abnova). Supernatants were diluted 1:5 in sample diluent and added to antibody-coated wells. Plates were incubated for 90 minutes at 37°C, washed with PBS, and incubated with biotinylated anti-galectin-1 detection antibody for 60 minutes at 37°C. After washing, avidin-biotin–horseradish peroxidase complex was added, followed by color development for 15 minutes at 37°C. Absorbance was measured at 450nm using a Cytation5 plate reader (BioTek).

### Imaging Mass Cytometry

Three indication kidney allograft biopsies with a diagnosis of ABMR, were included in the analysis. Serial sections were taken from formalin-fixed paraffin embedded kidney biopsies. One section was Periodic Acid Schiff-stained for the purpose of region of interest (ROI) selection. Remaining sections were 4µm-thick, mounted on positive charged slides. These slides were baked at 60℃ for 120 minutes prior to de-paraffinization using xylene, and incubations in descending grades of ethanol (100%, 95%, 80%, 70%). Antigen retrieval was conducted at 96℃ for 30 minutes (S236784-2; Agilent Technologies). Prior to antibody addition, tissues were covered with 3% BSA in PBS for 45 minutes in a hydration chamber. Incubation with all antibodies on the panel (**Supplemental Table S2**) also occurred in a hydration chamber, at 4℃ overnight. Slides were washed using 0.2% Triton X-100 in PBS, then stained with Ir-Intercalator for 30 minutes in the hydration chamber. Slides were air-dried and stored at room temperature prior to data acquisition on a Hyperion XTi imaging mass cytometer (Standard BioTools). Images for each ROI were acquired using the MCD Viewer software. This study was approved by the UHN Research Ethics Board CAPCR# 18-5489.

### Microfluidic Experiments

AngioPlate operational setup: AngioPlate devices were created in Dr. Boyang Zhang’s lab. The experimental procedure followed the previously published protocol (Rajasekar *et al*, 2025). Briefly, 25µL of a hydrogel mixture containing 10mg/mL fibrinogen (Sigma Aldrich, Cat# F3879) and 25U/mL thrombin (Sigma Aldrich, Cat# T6884) was added to each tissue well containing pre-patterned gelatin fibers. The fibrin hydrogel was allowed to crosslink at room temperature for 20 minutes. After gelation, the gelatin was degraded by adding 90µL of pre-warmed PBS supplemented with 1% aprotinin to each well. Following degradation, the PBS was removed, and a coating solution containing GMEC medium supplemented with 1 mg/mL fibrinogen, 0.1U/mL thrombin, and 1% aprotinin (2mg/mL, Sigma Aldrich, Cat# 616370-M) was added. The plate was then incubated overnight at 37°C. The following day, the coating solution was aspirated, and the plate was primed with GMEC medium containing 1% aprotinin for 48 hours prior to cell seeding. After priming, all media were aspirated, and 125µL of GMEC cell suspension (0.7 × 10⁶ cells/mL) was seeded into the inlet and outlet wells. Additionally, 30µL of GMEC medium was added to each tissue well. The plate was incubated statically at 37°C for 2 hours to allow GMECs to attach. Following cell attachment, perfusion was initiated using the IFlowRocker (OrganoBiotech, Cat# B001).

Endothelial barrier assessment in AngioPlate using dextran permeability assay: To assess the barrier integrity of glomerular microcapillaries in the AngioPlate, a 90µL mixture containing 1mg/mL of 4kDa FITC–dextran (Sigma-Aldrich, Cat#46944) and 65kDa TRITC–dextran (Sigma-Aldrich, Cat# T1162) was perfused through the inlet and outlet wells on day 7. These two sizes of dextran allow for detection of barrier integrity loss akin to glomerular filtration barrier damage, as damage can be detected by movement of 65kDa dextran, which is a similar size to the clinical assessment of albumin (62kDa) excretion. To maintain equal volumes across all wells, 65µL of culture medium was added to each tissue well. Fluorescence intensity readings were then measured, and the amount of dextran that leaked through the capillaries was quantified using standard curves generated from serial 1:10 dilutions of dextran.

Effect of IFNγ and *LGALS1* silencing on GMECs: To assess the effect of IFNγ on GMECs subjected to *LGALS1* knockdown or non-targeting control in the AngioPlate, microcapillaries were treated with 50U/mL of IFNγ. Vehicle control groups were also included, in which tissues were exposed to culture medium supplemented with 0.1% BSA in PBS. After 24 hours of treatment, endothelial barrier function was evaluated using the dextran permeability assay. Flowthrough from all wells were collected and analyzed for cytokine level quantification using a Human Pro-Inflammatory 15-Plex Assay (Eve Technologies, Cat# HDF15). Cytokines that were outside the assay’s detectable range were excluded from analysis. Following flowthrough collection, tissues were fixed and stained for CD31 (1:500; ab9498; Abcam), VE-cadherin (1:200; ab33168; Abcam), and galectin-1 (1:500; GTX101566; GeneTex), along with DAPI (Thermo Scientific). They were then incubated with the appropriate secondary antibody, either CF595-conjugated anti-rabbit secondary antibody (1:200; SAB4600107; Sigma) or FITC-conjugated anti-mouse secondary antibody (1:200; F0257; Sigma). Images were acquired using a Leica TCS SP8 confocal microscope (Leica Microsystems).

Effects of extracellular recombinant galectin-1 and *LGALS1* knockdown on GMECs: Recombinant galectin-1 (r-galectin-1) was produced and purified in Rabinovich’s lab as previously described (Barrionuevo *et al*, 2007). To assess the effect of r-galectin-1 on GMECs exposed to *LGALS1* knockdown or non-targeting control in the AngioPlate, microcapillaries were first treated with vehicle (0.1% BSA in PBS) or 50U/mL of IFNγ for 24 hours, followed by treatment with 10µM of r-galectin-1 for additional 24 hours. This dose of r-galectin-1 was chosen based on prior studies (Perillo *et al*, 1997; Nambiar *et al*, 2019; Toscano *et al*, 2007; Papa-Gobbi *et al*, 2021). After 24 hours of treatment, endothelial barrier function was evaluated using the dextran permeability assay. Flowthrough from all wells were collected and cytokine levels were measured as described above. One flowthrough sample was excluded from the analysis because it failed quality control.

### Statistical analysis

Graphical data are presented as mean ± SEM, Fold Change (FC), or Log2FC. For normally distributed data comparing two groups, Student’s t-test was used. For comparisons involving >2 groups with normally distributed data, one-way ANOVA with Tukey’s post hoc test was applied. Two-way ANOVA with Tukey’s post hoc test was used for normally distributed data with two separate treatments. For non-normally distributed data comparing two groups, the Mann-Whitney test was used. For non-normally distributed data comparing >2 groups with two separate treatments the Aligned Rank Transformation ANOVA was performed in R 4.2.3 using the package ARTool_0.11.2. Statistical significance was defined as p<0.05 or FDR<0.05.

### Bioinformatics

For differential proteins, pathway enrichment analysis was performed using pathDIP 5 (Pastrello *et al*, 2024) API in R. q-value 0.01 was used as a significance threshold, and categories “Drugs and vitamins” and “Disease” were excluded. Gene Ontology enrichment analysis was performed using Pantherdb (Thomas *et al*, 2022). Protein interactions among the differential proteins were identified using IID v.2021-05 (Kotlyar *et al*, 2026) and prioritized based on kidney annotation. Networks were built using NAViGaTOR 3.0.19.

## RESULTS

### GMECs are the main source of endogenous galectin-1 in kidney allografts with ABMR

We sought to identify cell types that express *LGALS1* gene and galectin-1 protein in healthy kidneys and kidneys with ABMR. To determine the cellular source of galectin-1 expression in the human kidney, we first analyzed publicly available single-cell RNA sequencing data (McEvoy *et al*, 2022). *LGALS1* expression in 19 healthy living donor kidney biopsies was predominantly in endothelial and immune cell subsets (**Figure 1A**). To compare galectin-1 expression between endothelial and immune cells, we performed flow cytometry on GMECs and PBMCs. Galectin-1 was robustly expressed in GMECs and minimally expressed in PBMCs (**Figure 1B**). Furthermore, immunofluorescence staining confirmed strong galectin-1 expression in both the cytoplasm and plasma membrane of primary human GMECs (**Figure 1C-D**).

**Figure 1.**
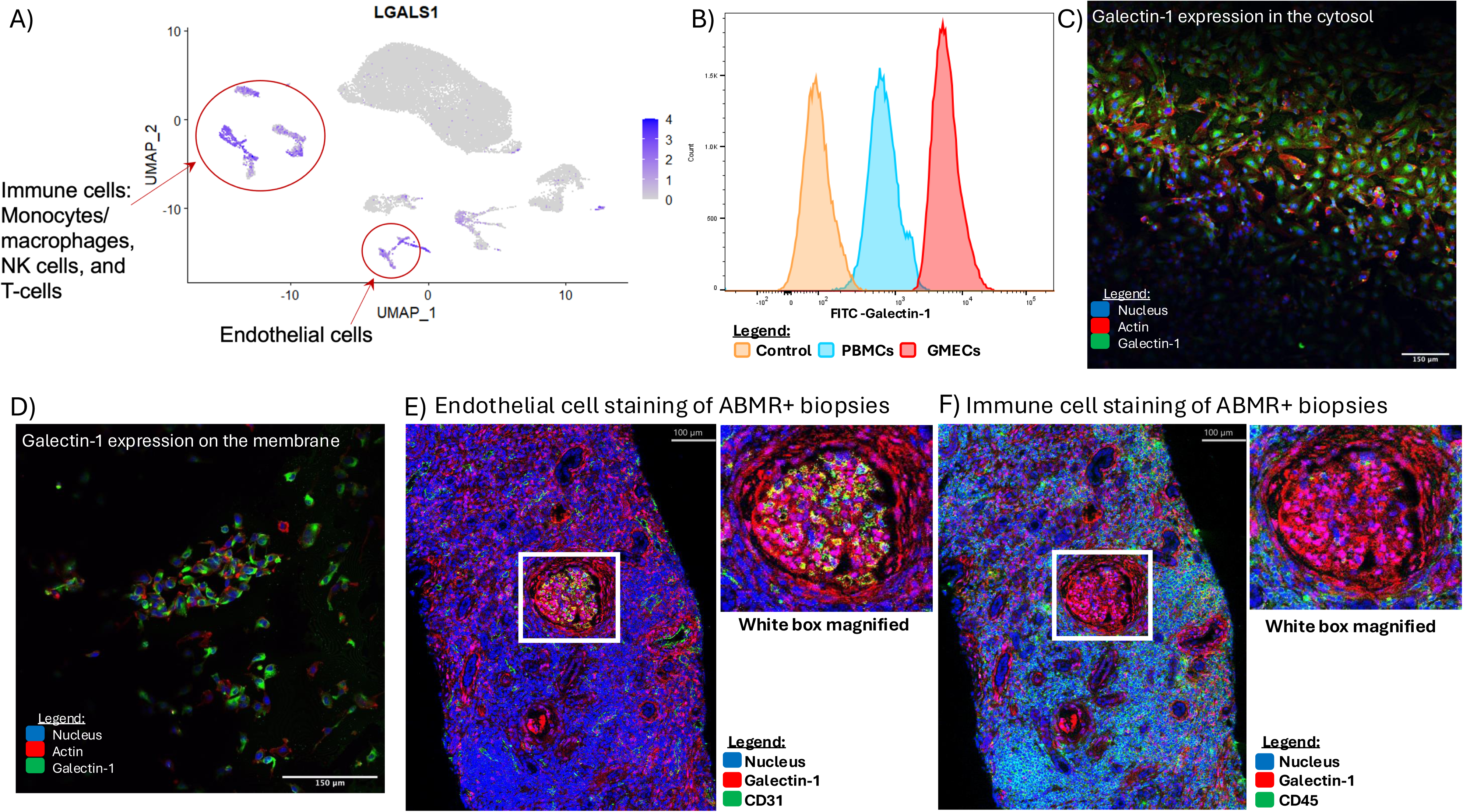
Glomerular microvascular endothelial cells (GMECs) are the main cell expressing galectin-1 in the kidney. **A)** Feature plot showing *LGALS1* gene expression in immune cells and endothelial cells (purple) and no expression in other cell types (grey), in the healthy human kidney, based on single-cell RNA-Sequencing data from 19 living donor biopsies (McEvoy *et al*, 2022). **B)** Expression of galectin-1 in GMECs vs PBMCs by flow cytometry. Galectin-1 Absent Control (orange), PBMC (blue), GMEC (red). **C)** Galectin-1 protein expression in the cytoplasm and **D)** on the membrane of primary GMECs stained for galectin-1 (green), DAPI (blue) and actin (red). **E)** Imaging mass cytometry comparing CD31 and galectin-1 co-expression to **F)** CD45 and galectin-1 co-expression in a representative kidney biopsy from a patient with ABMR. White boxes magnify the glomerulus. CD31 or CD45 (green), nuclei (blue) and galectin-1 (red). NK cells, Natural Killer cells; GMEC, Glomerular Microvascular Endothelial Cells; FMO, Fluorescence minus one; PBMC, Peripheral Blood Mononuclear Cell; ABMR, Antibody Mediated Rejection.

We next used imaging mass cytometry (IMC) to determine the sites of galectin-1 expression in kidney allograft biopsies from patients with ABMR. In ABMR biopsies, galectin-1 co-localized extensively with CD31-positive endothelial cells (**Figure 1E**) but showed minimal co-staining with CD45-positive immune cells in the glomerulus (**Figure 1F**). This finding was consistent in other biopsies from patients with ABMR and healthy controls (**Supplemental Figure S4 and S5)**. Together, these findings indicate that GMECs are the predominant source of galectin-1 in ABMR.

### *LGALS1* knockdown does not alter GMEC phenotype *in vitro*

We next designed a study to test the role of endogenous galectin-1 on the cell phenotype, proteomic profile, and function of primary human GMECs *in vitro*, in the presence or absence of IFNγ (**Figure 2**). Electroporation was used to deliver siRNA targeting *LGALS1*, and RT-qPCR confirmed a knockdown efficiency of 80%, 48 hours post-electroporation (**Supplemental Figure S6A**). At 72 hours post-electroporation, the levels of galectin-1 protein secreted by these GMECs were also significantly decreased (P<0.001) (**Supplemental Figure S6B**). Next, we assessed expression of the phenotypic GMEC markers *PECAM1* (CD31), *CDH5* and *VWF* at 48, 72, and 96 hours post-electroporation. None of these genes changed significantly at any of the time points analyzed (**Supplemental Figure S6C**). Moreover, we assessed expression of GMEC activation markers *ACTA2*, *IL6*, and *ICAM1* (**Supplemental Figure S6D**). Again, none of these transcripts changed significantly at any of the time points analyzed. Collectively, these findings demonstrate that *LGALS1* knockdown does not perturb baseline GMEC phenotype or activation, allowing for further downstream proteomic analyses.

**Figure 2.**
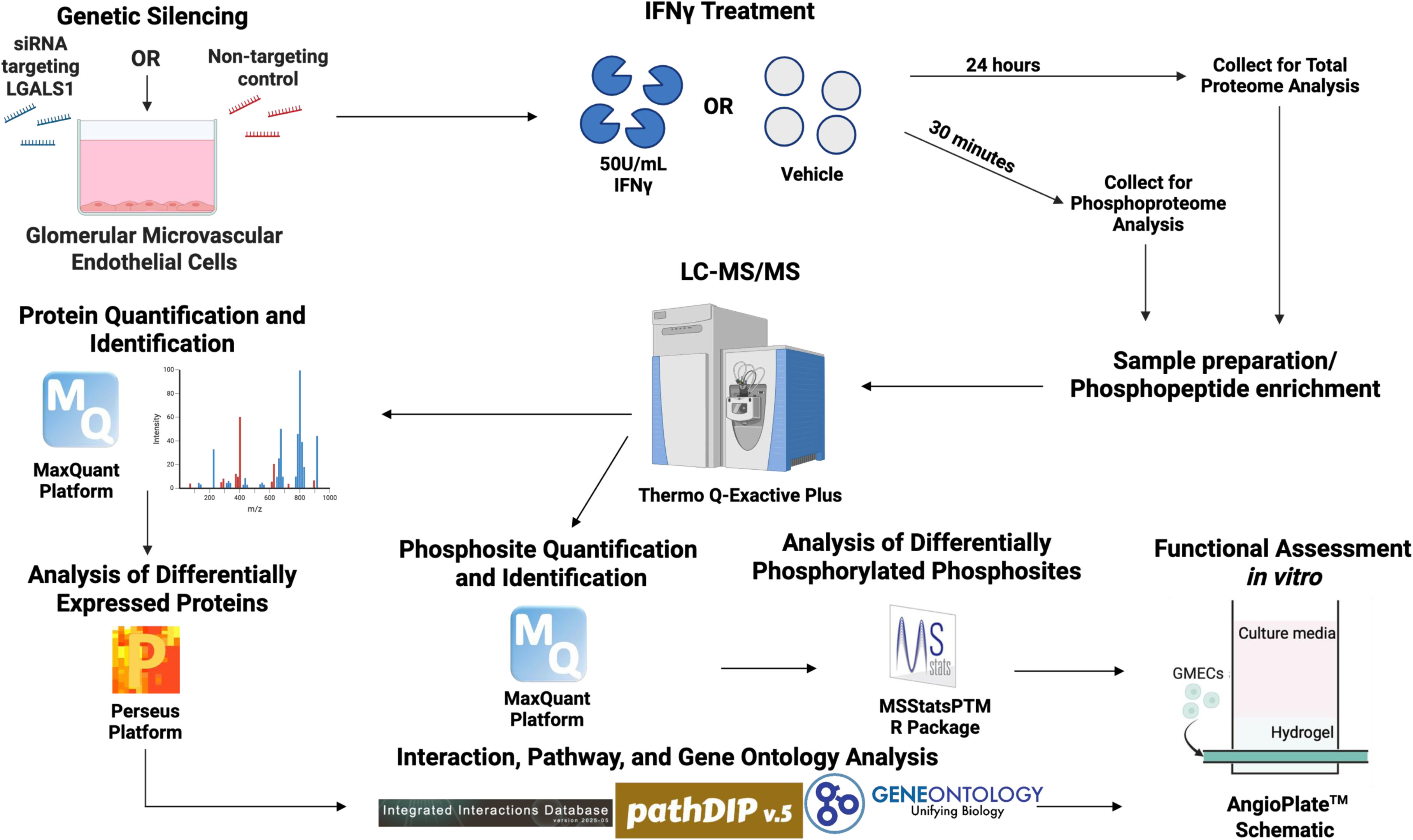
Proteomics and phosphoproteomics workflow. Overview of proteomics/phosphoproteomics workflow, including genetic inhibition of *LGALS1* in glomerular microvascular endothelial cells (GMECs), treatment of GMECs with 50U/mL of interferon-γ (IFNγ) or vehicle, sample preparation and phosphopeptide enrichment, followed by LC-MS/MS, data analysis by MaxQuant, statistical analysis to identify differentially expressed proteins by Perseus for total proteomics, or MSStatsPTM for phosphoproteomics, pathway enrichment analysis (PathDIP) (Pastrello *et al*, 2024), protein-protein interaction analysis (IID) (Kotlyar *et al*, 2026), and Gene Ontology enrichment analysis (Ashburner *et al*, 2000; Thomas *et al*, 2022; The Gene Ontology Consortium, 2026), followed by functional assessment in AngioPlate glomerular capillary *in vitro*. siRNA, small interfering ribonucleic acid; *LGALS1*, galectin-1 gene; IFNγ, interferon-γ; LC-MS/MS, liquid chromatography followed by tandem mass spectrometry; PTM, post-translational modification.

### Changes in the immune system and ECM proteins in IFNγ-treated GMECs

We next sought to determine the impact of *LGALS1* silencing and IFNγ treatment on the GMEC proteome (**Figure 3A**). First, we confirmed that galectin-1 protein was significantly decreased (by >2.8 fold) in GMECs subjected to *LGALS1* knockdown (**Figure 3B**). We then performed an unbiased proteomics analysis and identified 5446 proteins in lysates of *LGALS1*-silenced GMECs treated with IFNγ or vehicle control (**Figure 3C**). After filtering, imputation, and batch correction, we performed statistical analysis on 5186 proteins. We identified 827 proteins that were significantly differentially expressed in response to IFNγ treatment compared to vehicle control (2-way ANOVA and Tukey’s post-test, FDR<0.05) (**Figure 3C**). Unsupervised hierarchical clustering of these differentially expressed proteins following IFNγ treatment revealed two distinct clusters corresponding to proteins with increased or decreased abundance (**Supplemental Table S3**). Treatment with IFNγ was characterized by up-regulation of pathways related to immune system and interferon signaling (**Supplemental Figure S7A, Supplemental Table S4)**, and downregulation of pathways associated with ECM organization and cell–cell interactions (**Supplemental Figure S7B, Supplemental Table S5)**. These proteomic changes are consistent with prior *in vitro* observations at both transcript and protein levels (Clotet-Freixas *et al*, 2020; Diaz & Jiménez, 1997; Serpier *et al*, 1997), supporting the robustness of the IFNγ-induced molecular programs observed in this system.

**Figure 3.**
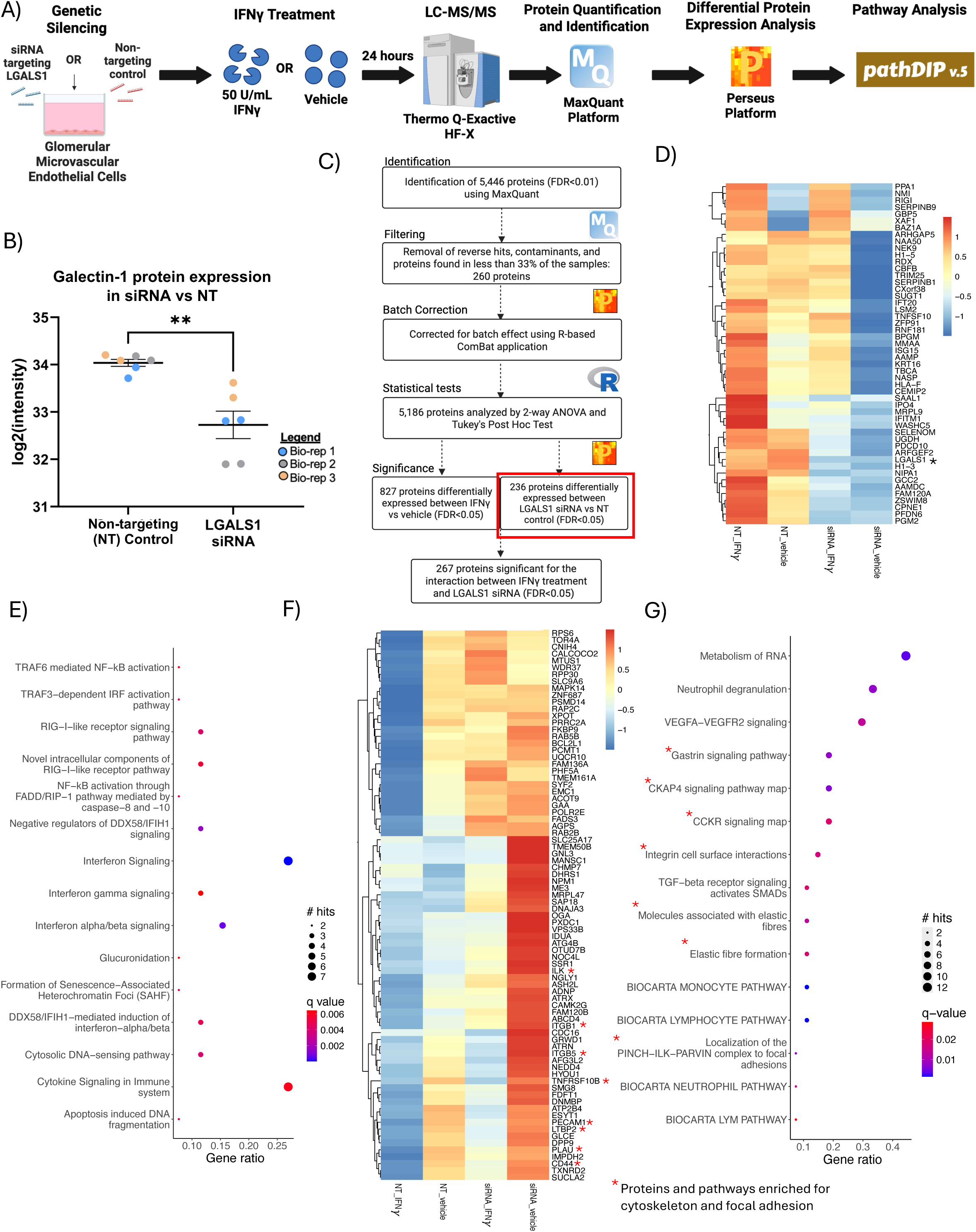
*LGALS1* silencing in human GMECs controls expression of ECM and immune system proteins. **A)** Overview of GMEC treatment and proteome analysis workflow. **B)** Galectin-1 protein expression was significantly decreased with *LGALS1* siRNA treatment. Data are presented as mean ± SEM. Student’s T-test, **P<0.01. **C)** Proteins identified, quantified, and significant for each treatment and their interaction. Red box denotes proteins significant with *LGALS1* siRNA alone. **D)** Heatmap of 50 proteins included in cluster 3 (complete heatmap is in **Supplemental Figure S6**). Red colour indicates increased expression and blue colour indicates decreased expression, with asterisk denoting galectin-1. **E)** Bubble plot of top 15 pathways enriched among cluster 3 proteins (Benjamini Hochberg, FDR<0.01). The size of the circle indicates the number of proteins in the pathway, and the colour indicates significance. Pathways extracted from PathDIP (Pastrello *et al*, 2024). **F)** Heatmap of 80 proteins included in cluster 1 (complete heatmap is in **Supplemental Figure S7**). Red asterisk denotes proteins mapped to cytoskeleton and focal adhesion pathways. **G)** Bubble plot of top 15 pathways enriched among cluster 1 proteins (Benjamini Hochberg, FDR<0.01). ECM and focal adhesion pathways are denoted by red asterisk. Pathways extracted from PathDIP (Pastrello *et al*, 2024) *LGALS1*, galectin-1 gene; GMEC, glomerular microvascular endothelial cells; NT, non-targeting control; Bio-rep, biological replicate; ECM, extracellular matrix.

### *LGALS1* silencing increases expression of ECM and focal adhesion proteins and decreases IFNγ and immune response-related proteins

To determine the role of endogenous galectin-1 in modulation of protein expression in glomerular endothelial cells, we next examined the proteome of GMECs altered in response to *LGALS1* knockdown. A total of 236 proteins were found to be differentially expressed in response to *LGALS1* silencing compared to non-targeting control (**Figure 3C**) and plotted in a heatmap using unsupervised hierarchical clustering. The heatmap was divided into 4 clusters based on differential expression using the second layer of the hierarchy (**Supplemental Figure S8**). Of particular interest were clusters 1 and 3. The 50 proteins in cluster 3 had decreased expression with *LGALS1* knockdown and unsurprisingly, contained galectin-1 protein as well (**Figure 3D**). Interestingly, upon pathway analysis (Pastrello *et al*, 2024), these proteins were enriched for cytokine and IFN signaling pathways, suggesting that inhibiting *LGALS1* resulted in the opposite effects to IFNγ stimulation (**Figure 3E, Supplemental Table S6**). In contrast, the 80 proteins in cluster 1 had increased expression following *LGALS1* knockdown (**Figure 3F)**. Some of these proteins included CD44, PECAM1 (CD31), PLAU, ITGB1, and ITGB5, which are instrumental for endothelial phenotype and adhesion to the ECM (Fernández-Martín *et al*, 2012). Proteins in cluster 1 demonstrated 19 enriched pathways (hypergeometric test, FDR: BH<0.01) (**Supplemental Table S7**), including pathways involving ECM, focal adhesion, and immune cell interaction, such as molecules associated with elastic fibres (q= 5.39E-04), elastic fibre formation (q= 9.00E-04), integrin cell surface interaction (q= 4.94E-04), and localization of the PINCH-ILK-PARVIN complex to focal adhesions (q=1.10E-04) (**Figure 3G**). Loss of galectin-1 caused proteomic changes involving IFN-associated immune pathways, ECM, and adhesion-related programs in GMECs.

### IFNγ stimulation and *LGALS1* silencing synergize to alter expression of focal adhesion and cytoskeletal proteins in GMECs

To further dissect the combined effects of IFNγ treatment and *LGALS1* knockdown on endothelial cell regulation, we identified proteins with statistically significant interaction effects between the two treatments in GMECs. In total, 267 proteins were significantly altered due to the interaction between IFNγ stimulation and *LGALS1* silencing (**Figure 4A**). We applied unsupervised hierarchical clustering on these 267 proteins (**Supplemental Figure S9**) and performed Gene Ontology (GO) enrichment analysis (Ashburner *et al*, 2000; Thomas *et al*, 2022; The Gene Ontology Consortium, 2026) to investigate if concurrent IFNγ stimulation and *LGALS1* knockdown alter similar immune and ECM processes as the individual treatments. GO enrichment analysis revealed that these proteins were highly enriched in biological processes related to extracellular matrix organization and cell adhesion (full list of enriched GO terms is included as **Supplemental Tables S8-10**). Of particular interest among the top 20 enriched terms were regulation of cytoskeleton (q=4.58E-03, GO-Biological Process) **(Figure 4B**), focal adhesion (q=5.25E-04, GO-Cellular Component), anchoring junction (q=1.14E-03, GO-Cellular Component) **(Figure 4C**), and cell adhesion molecule binding (q=2.41E-02, GO-Molecular Function) (**Figure 4D**). These terms are all related to cytoskeleton and focal adhesion, processes important for the formation and function of the glomerular filtration barrier, which is damaged during chronic ABMR (Chutani *et al*, 2024).

**Figure 4.**
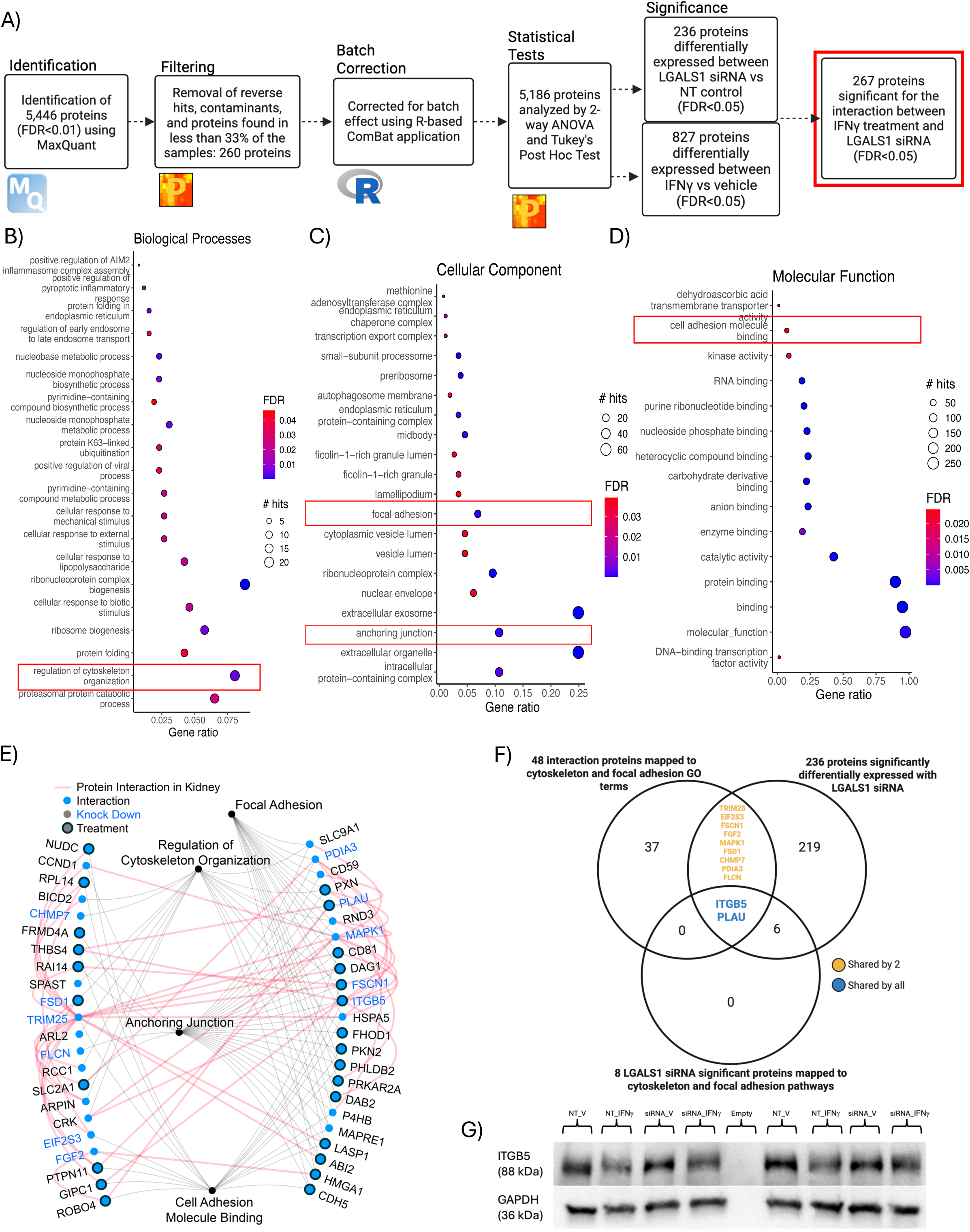
IFNγ and *LGALS1* silencing alter expression of focal adhesion proteins in GMECs. **A)** Proteins identified, quantified, and significant for each treatment and their interaction; red box denotes proteins significant for the interaction between IFNγ treatment and *LGALS1* siRNA. **B-D)** Bubble plots of top 20 Gene Ontology (GO) terms (Biological Process, Cellular Component, and Molecular Function) enriched among the 267 interaction-significant proteins (FDR: BH<0.01). The size of the circle indicates the number of proteins in the pathway, and the colour indicates significance. Cytoskeleton and focal adhesion terms are denoted by red boxes. **E)** Protein-protein interaction network (generated using IID (Kotlyar *et al*, 2026)) of significant proteins mapped to focal adhesion, regulation of cytoskeleton organization, anchoring junction, and cell adhesion molecule binding GO terms. The connection between the nodes and mapped GO terms are shown in grey. Physical protein interactions annotated with kidney expression are shown in red. Nodes represented with a blue dot are significant for the interaction between *LGALS1* siRNA and IFNγ. Nodes with an outside circle are significant for IFNγ vs vehicle treatment. Nodes with protein names in blue are significant for *LGALS1* siRNA vs NT control. **F)** Nine proteins were shared between the 48 significant interaction proteins mapped to GO terms and the 236 proteins significantly altered by *LGALS1* siRNA. The two proteins, ITGB5 (Integrin subunit beta 5) and PLAU (Urokinase-type plasminogen activator), were shared by all 3 datasets, including the 2 previously mentioned and the 8 proteins significantly altered by *LGALS1* siRNA, and mapped to cytoskeleton and focal adhesion pathways from Figure 3. **G)** Validation of β5 integrin (ITGB5) expression changes using immunoblotting. GAPDH (Glyceraldehyde 3-phosphate dehydrogenase) was included as a loading control. *LGALS1*, galectin-1 gene; GMEC, glomerular microvascular endothelial cells; IFNγ, interferon-γ; GO, Gene Ontology; NT, non-targeting control.

To further contextualize these findings, the Integrated Interactions Database (Kotlyar *et al*, 2026), was searched for experimentally validated interactions among the 48 proteins mapped to the GO terms highlighted above, revealing a highly interconnected protein network (**Figure 4E**). Several proteins within this network, such as CDH5 (VE-cadherin), CD81, ROBO4, and FASCN1, independently reached significance for IFNγ treatment or *LGALS1* knockdown, indicating convergent regulation of focal adhesion pathways by inflammatory signaling and galectin-1. Comparison of *LGALS1* knockdown/IFNγ interaction significant proteins with those altered by *LGALS1* knockdown alone identified a subset of overlapping focal adhesion–related proteins, including ITGB5 and PLAU, linking galectin-1-dependent regulation to IFNγ-modulated cytoskeletal pathways (**Figure 4F**). Finally, differential expression of ITGB5 due to IFNγ and *LGALS1* knockdown was validated by immunoblotting to confirm proteomic findings observed in mass spectrometry data (**Figure 4G, Supplemental Figure S10**). Collectively, these analyses identified focal adhesion and ECM proteins as major targets of the interaction between IFNγ treatment and *LGALS1* knockdown in GMECs.

### *LGALS1* silencing and IFNγ treatment of GMECs alter the phosphorylation of proteins involved in cytoskeleton organization, endocytosis, and caveolae formation

To investigate the mechanisms through which IFNγ treatment and *LGALS1* silencing affect signaling, we performed phosphoproteomic profiling of treated GMECs. An overview of the treatment groups and phosphopeptide enrichment workflow is shown in **Figure 5A**. Using label-free quantification, we identified 4,140 unique proteins (FDR<0.01) and 2,727 unique phosphopeptides (FDR<0.01) (**Figure 5B).** The MSstatsPTM package in R was utilized to quantify phosphopeptides across all conditions and 28 phosphopeptides were significantly altered (p<0.01) after correcting for the abundance of their matched unmodified peptides (**Figure 5B**, **Supplemental Table S11**). Volcano plots summarizing differential phosphorylation revealed distinct phosphosite signatures associated with IFNγ treatment, *LGALS1* knockdown, and their interaction (**Figure 5C**). Interestingly, across all 3 comparisons the same phosphopeptide of caveolae-associated protein 1 (CAVN1), Q6NZI2_S169_S171 was significantly decreased in response to IFNγ treatment, *LGALS1* knockdown, and their interaction (FDR<0.2). In addition, phosphopeptide Q09666_S5752 (AHNK protein) was significantly increased by both IFNγ treatment and *LGALS1* knockdown (FDR<0.2). The phosphopeptide Q07157_S166_S168 (ZO1 protein) was also decreased by IFNγ treatment but failed to reach statistical significance after adjustment. We next focused on 28 phosphopeptides that were significantly and differentially phosphorylated in at least one treatment condition. We generated a supervised heatmap illustrating phosphopeptide expression across conditions, and their enriched functional categories (**Figure 5D**). Notably, several of the phosphosites, including ZO1, AHNK, and CAVN1, mapped to proteins involved in cell adhesion, cytoskeleton, and endocytosis.

**Figure 5.**
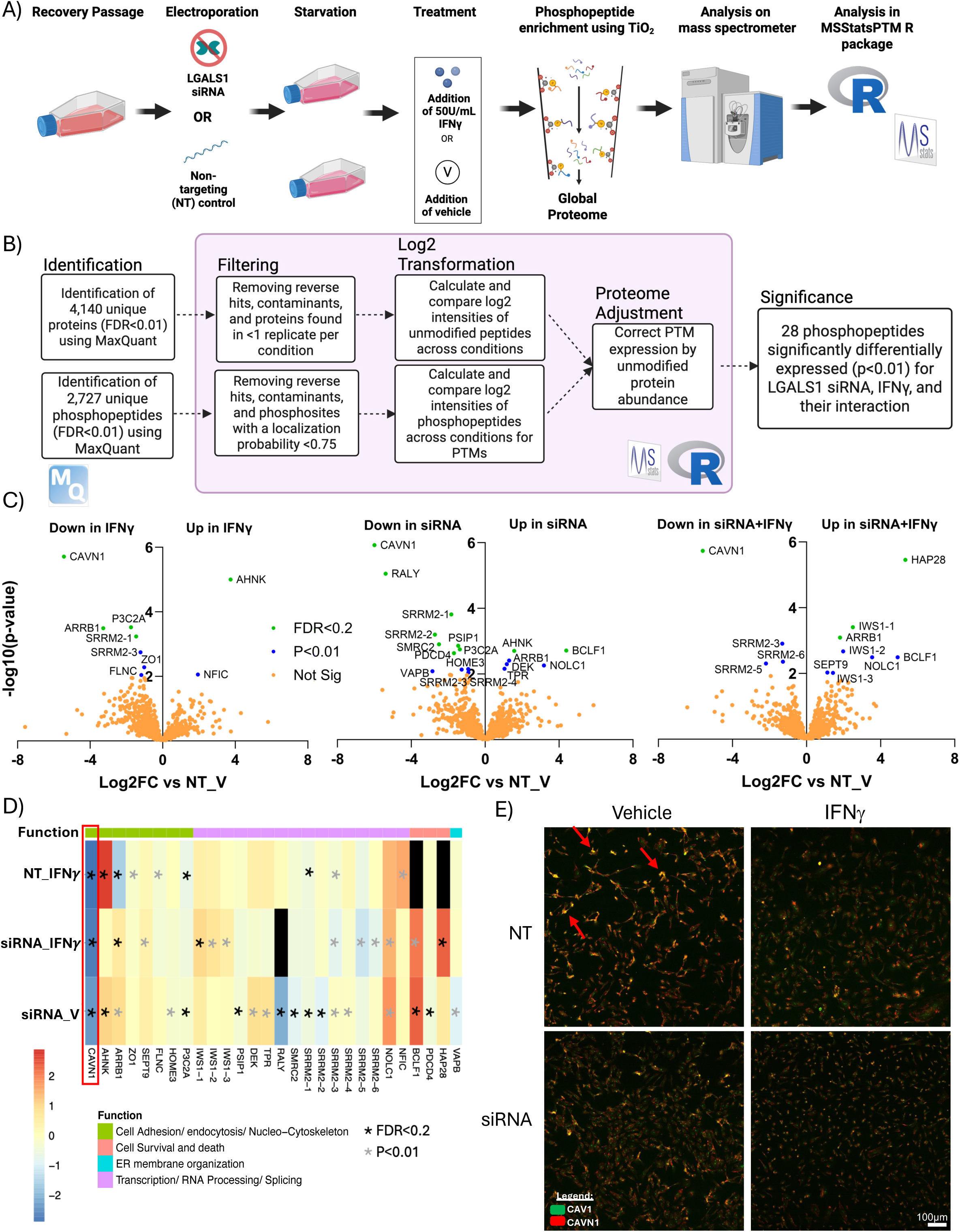
*LGALS1* silencing together with IFNγ treatment alter the phosphorylation of proteins involved in cytoskeleton organization, endocytosis and caveolae formation in GMECs. **A)** Overview of treatment and phosphoproteome enrichment workflow. **B)** Phosphopeptides identified, quantified, and significant for each treatment and their interaction after correcting for matched unmodified peptides using MSStatsPTM in R. **C)** Volcano plots of phosphopeptide Log2FC of IFNγ treatment, *LGALS1* siRNA, and both IFNγ and *LGALS1* siRNA, compared to non-targeting control and vehicle plotted against –log10(p-value). Orange colour represents non-significant proteins, while blue colour represents those with p<0.01 (linear mixed-effects model), and green represents those with FDR: BH<0.2. **D)** Heatmap of 28 phosphopeptides significant in at least 1 comparison from panel C. Red colour indicates increased expression, blue colour indicates decreased expression, and black colour indicates phosphopeptides that were not found in that comparison. Colour on the top bar denotes function (obtained from uniport.org) of the protein the phosphopeptide belongs to. Black and grey asterisks denote FDR<0.2 and p<0.01, respectively. Immunostaining of CAVN1 and CAV1 when GMECs were treated with *LGALS1* siRNA, IFNγ, or both. Green colour indicates CAV1, red colour indicates CAVN1. Red arrows denote co-staining of CAV1 and CAVN1 in the NT_V condition. *LGALS1*, galectin-1 gene; GMEC, glomerular microvascular endothelial cells; IFNγ, interferon-γ; NT, non-targeting control; NT_V, non-targeting control and vehicle; PTM, post-translational modification.

We further investigated the effect of decreased CAVN1 phosphorylation in GMECs due to its consistent changes across conditions and existing literature showing that CAVN1 phosphorylation is critical for recruitment of caveolin-1 (CAV1) to caveolae, flask-like structures that mediate mechanosensing and endocytosis, and are increased in chronic ABMR (Hill *et al*, 2008; Yamanaka *et al*, 2019; Teixeira *et al*, 2021). Further prior evidence also suggested that CAVN1 can be dissociated from CAV1 in the context of cellular perturbation such as osmotic stress or cellular senescence and enter the nucleus to mediate signaling (Bai *et al*, 2011; Qifti *et al*, 2022). To investigate the possibility that reduced CAVN1 phosphorylation in response to IFNγ treatment or *LGALS1* knockdown results in dissociation between CAVN1 and CAV1, we next performed immunofluorescence to visualize CAVN1 and CAV1 co-expression in GMECs. While CAVN1 and CAV1 colocalized in vehicle-treated GMECs, their colocalization was diminished in IFNγ-treated or *LGALS1* knockdown cells (**Figure 5E, Supplemental Figure S11**). This loss of interaction between CAVN1 and CAV1 suggests a disruption in caveolar assembly and membrane anchoring in response to galectin-1 depletion and cytokine stimulation.

### Manipulation of galectin-1 expression levels alters GMEC vascular permeability in conjunction with IFNγ treatment

To model glomerular microvascular responses relevant to ABMR, we used a microfluidic platform called AngioPlate (Rajasekar *et al*, 2025). AngioPlate is a customized 384-well plate in which three adjacent wells constitute a single, independent perfusable unit (**Figure 6A**), allowing for multiplexed assessment of the effect of *LGALS1* siRNA, IFNγ treatment, and the addition of r-galectin-1. Each unit contained a pre-patterned gelatin fiber that is degraded to generate a hollow, perfusable channel embedded within a natural hydrogel matrix. These channels were subsequently seeded with primary GMECs, resulting in the formation of confluent, 3D perfusable glomerular microcapillaries.

**Figure 6.**
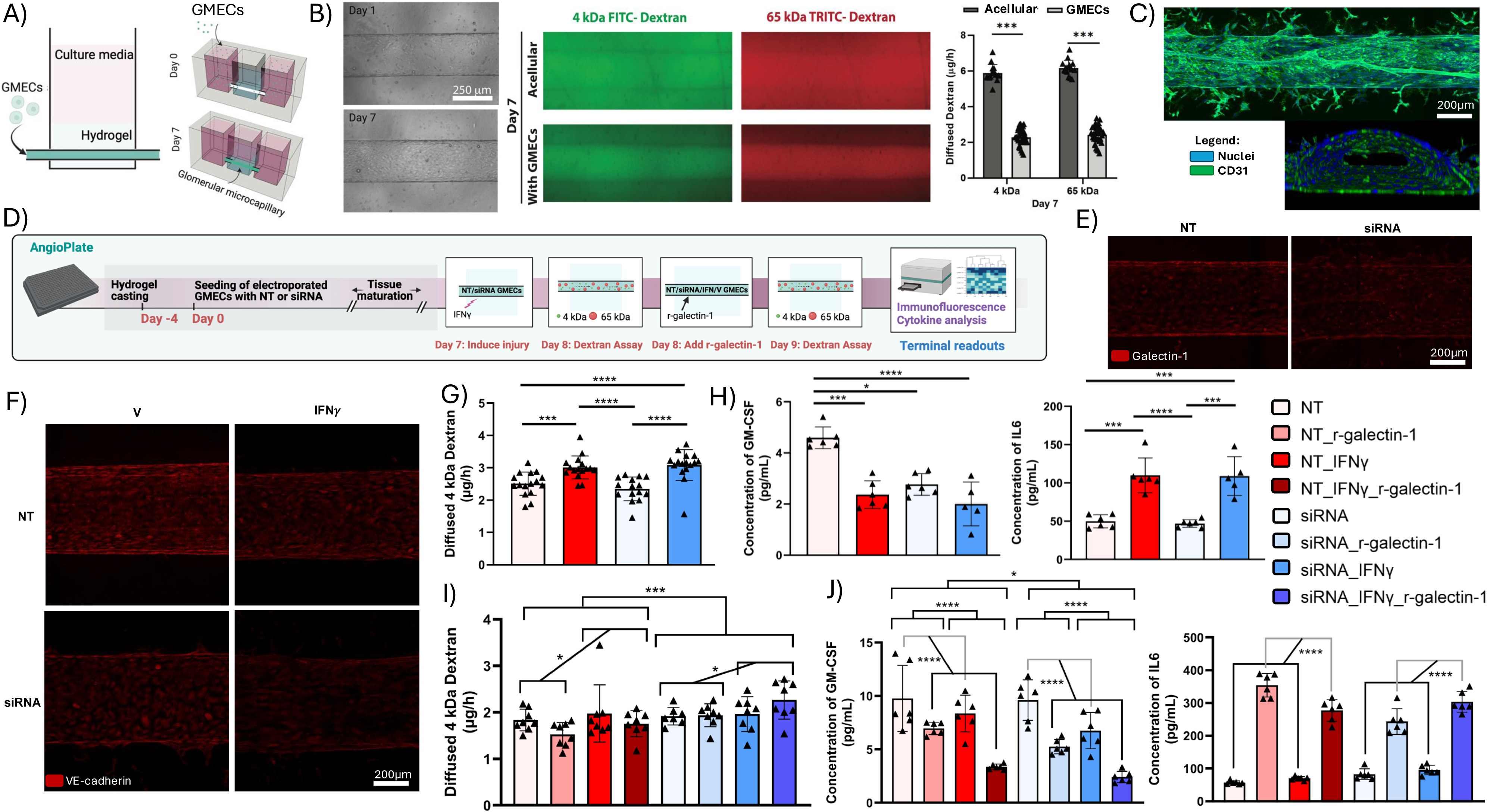
*LGALS1* silencing, recombinant galectin-1, and IFNγ treatment alter GMEC vascular permeability and cytokine secretion. **A)** Schematic diagram of AngioPlate™ preparation and cell seeding. **B)** Capillaries containing confluent GMECs in AngioPlate™ form tight barrier to both 4kDa and 65kDa dextran on day 7 after seeding, compared to acellular tubes. Data presented as mean ±SEM, n=16-32 per group, Holm-Sidak, ***P<0.001. **C)** Immunofluorescence image of microvessel stained with CD31 (green) and DAPI (blue). **D)** Overview of AngioPlate™ timeline including setup, seeding of GMECs pretreated with NT or *LGALS1* siRNA, permeability assays post-IFNγ and r-galectin-1 treatments, and the terminal cytokine measurement in the flowthrough. **E)** Representative immunofluorescence image of the microvessel comparing expression of galectin-1 (red) in *LGALS1* siRNA to non-targeting (NT) control. **F)** Representative immunofluorescence image of the microvessel comparing expression of VE-cadherin (red) in *LGALS1* siRNA or NT control, and IFNγ treatment or vehicle. **G)** GMEC vessel barrier integrity was assessed by diffusion of 4kDa dextran in µg/hour after IFNγ treatment and/or *LGALS1* siRNA treatment. Data presented as mean ±SEM, n=15-16 per group, Aligned Rank Sum Test, ***P<0.001 ****P<0.0001. **H)** GM-CSF and IL-6 cytokine concentrations in flowthrough from wells treated with *LGALS1* siRNA, IFNγ, both, or controls. Data presented as mean ±SEM, n=5-6 per group, Aligned Rank Sum Test, *P<0.05, ***P<0.001, ****P<0.0001. **I)** GMEC vessel barrier integrity was assessed by diffusion of 4kDa dextran in µg/hour after IFNγ Treatment and/or *LGALS1* siRNA treatment, followed by the addition of r-galectin-1 or vehicle. Data presented as mean ±SEM, n=7-8 per group, Aligned Rank Sum Test, *P<0.05, ***P<0.001. **J)** GM-CSF and IL-6 cytokine concentration in flowthrough from wells treated with *LGALS1* siRNA, IFNγ, both, or control, followed by the addition of r-galectin-1 or vehicle. Data presented as mean ±SEM, n = 5-6 per group, Aligned Rank Sum Test, *P<0.05, ***P<0.001, ****P<0.0001. GMECs, glomerular microvascular endothelial cells; FITC-dextran, fluorescein conjugated dextran; TRITC-dextran, Tetramethylrhodamine conjugated dextran; CD31, *PECAM1* gene; NT, non-targeting control and vehicle; NT_r-galectin-1, non-targeting control and vehicle and recombinant galectin-1; NT_IFNγ, non-targeting control and IFNγ treatment; NT_IFNγ_r-galectin-1, non-targeting control and IFNγ and recombinant galectin-1; siRNA, *LGALS1* siRNA and vehicle; siRNA_r-galectin-1, *LGALS1* siRNA and vehicle and recombinant galectin-1; siRNA_IFNγ, *LGALS1* siRNA and IFNγ treatment; siRNA_IFNγ_r-galectin-1, *LGALS1* siRNA and IFNγ and recombinant galectin-1.

By day 7, GMEC-lined channels exhibited a significantly tighter barrier to both 4 kDa and 65 kDa dextran polymer compared with acellular controls, indicating the establishment of an active endothelial barrier (**Figure 6B**). Further characterization confirmed uniform endothelial coverage throughout the channel, with robust expression of CD31, a key endothelial junctional marker (**Figure 6C**). Using this system, we evaluated how IFNγ and modulation of *LGALS1* influence vascular integrity and cytokine production (**Figure 6D)**. *LGALS1* knockdown resulted in decreased galectin-1 protein expression in the vessel (**Figure 6E)**. Silencing *LGALS1* in conjunction with IFNγ treatment showed a decrease in VE-cadherin expression, replicating results from the proteome data (**Figure 6F)**. We next examined vascular permeability after distinct treatments. Exposure to IFNγ led to a significant loss of barrier integrity (p=3.06E-11), reflected by increased diffusion of both 4kDa (**Figure 6G)** and 65kDa dextran (**Supplemental Figure 12A)**. Every treatment that was added to the cells had a more pronounced effect on 4kDa dextran than 65kDa, as expected with the difference in dextran size. Silencing of *LGALS1* alone did not show any effect on the barrier; however, the interaction of *LGALS1* silencing in tandem with IFNγ treatment had a significant effect (Aligned Rank-Sum Test, 2-way ANOVA, *LGALS1* siRNA x IFNγ treatment p=0.039) on increasing 4kDa diffusion (**Figure 6G)** and no effect on 65kDa dextran diffusion (**Supplemental Figure 12A)**. Remarkably, *LGALS1* knockdown combined with IFNγ treatment increased permeability and reduced VE-cadherin expression throughout the vessel, consistent with impaired junctional organization. We next evaluated cytokine secretion by the endothelium after perturbation. The vessel flowthrough media demonstrated a significant reduction in GM-CSF secretion by GMECs in response to IFNγ treatment, *LGALS1* knockdown, or their combination (**Figure 6H**). Interestingly, GM-CSF was previously shown to promote angiogenesis, endothelial cell adhesions and microvessel density (Zhao *et al*, 2014; Yan *et al*, 2017), suggesting that this decrease in response to IFNγ treatment and *LGALS1* silencing may be responsible for reduced vessel competence. In contrast, IL-6 levels increased significantly only in response to IFNγ (**Figure 6H)**. Similar increases in MCP-1 were observed with IFNγ treatment, and elevated IFNγ levels were verified in microvessels exposed to this cytokine (**Supplemental Figure 12B**).

We next examined the effects of r-galectin-1 on microvessel permeability and endothelial function. Notably, the effects of r-galectin-1 were dependent on prior *LGALS1* silencing, as r-galectin-1 had a distinct effect on permeability depending on whether endogenous *LGALS1* was silenced. The addition of r-galectin-1 induced increased permeability with both *LGALS1* knockdown and IFNγ, but decreased permeability in the absence of these two treatments, with both 4kDa (**Figure 6I)** and 65kDa dextran (**Supplemental Figure 12C**). Notably, r-galectin-1 also decreased GM-CSF concentrations in the GMEC secretome, to a greater degree than either IFNγ or *LGALS1* silencing, suggesting an interplay between r-galectin-1 and both IFNγ treatment and *LGALS1* silencing (**Figure 6J)**. In contrast, r-galectin-1 dramatically increased IL-6 levels, to levels higher compared to IFNγ stimulation (**Figure 6J)**. Interestingly, r-galectin-1 significantly increased other proinflammatory cytokines, including IL-8 and MCP-1, and promoted an increase in IFNγ above the levels induced by IFNγ stimulation (**Supplemental Figure S12D**). These findings indicate that IFNγ increases GMEC permeability, possibly by reducing VE-cadherin expression and GM-CSF secretion. While *LGALS1* knockdown alone had minimal effects on permeability, it significantly altered GM-CSF secretion and reversed the impact of the exogenous r-galectin-1. In contrast, r-galectin-1 decreased permeability under baseline conditions but promoted increased permeability in the presence of IFNγ or *LGALS1* knockdown. Additionally, r-galectin-1 markedly enhanced proinflammatory cytokine secretion, highlighting a key role for this lectin in regulating GMEC function and phenotype.

## DISCUSSION

Motivated by the incomplete understanding of ECM remodeling in chronic ABMR and other glomerular diseases, and our prior observation that galectin-1 was increased in kidney glomeruli with ABMR, coinciding with decreased ECM proteins, this study was conducted to dissect the effects of galectin-1 manipulation in primary human GMECs. Our study revealed three main observations. First, galectin-1 was predominantly expressed in GMECs in biopsies with ABMR. Second, *LGALS1* knockdown, in tandem with IFNγ stimulation, resulted in perturbation of cytoskeletal and adhesion proteins, and disruption of caveolar proteins in GMECs. Third, using an innovative microfluidic platform to mimic a glomerular capillary, we showed that extracellular r-galectin-1 decreased vessel permeability when administered alone, although it significantly increased permeability in the presence of IFNγ or *LGALS1* silencing, likely due to altered expression of adhesion and ECM proteins. This study expands the knowledge of galectin-1 biology in primary GMECs and sets the stage for future *in vivo* examination of its role as a therapeutic target in ABMR.

We established GMECs as the predominant cells expressing galectin-1 within the glomerulus. The endothelial cell origin is consistent with previous reports implicating galectin-1 in endothelial cell migration, angiogenesis, and tissue repair, and in fibrotic remodeling when repair processes become dysregulated (Griffioen & Thijssen, 2014). In the context of ABMR, where endothelial cells are chronically exposed to proinflammatory stimuli such as donor-specific antibodies and cytokines like TNFα and IFNγ (Lion *et al*, 2016), galectin-1 may therefore act as a critical regulator of endothelial adaptive versus maladaptive responses.

Proteomic analysis following *LGALS1* silencing revealed coordinated alterations in ECM, focal adhesion, and immune signaling pathways without perturbing baseline endothelial identity or viability. Notably, *LGALS1* depletion resulted in reduced expression of IFN-associated immune response proteins alongside increased abundance of adhesion- and matrix-associated proteins. This reciprocal regulation suggests that endogenous galectin-1 functions as a molecular switch balancing inflammatory signaling with structural and adhesive programs in GMECs. Such a role is particularly relevant to kidney allograft rejection, where inflammation coexists with progressive glomerular basement membrane thickening in chronic ABMR (De Serres *et al*, 2011). Several adhesion-related proteins altered by *LGALS1* knockdown have well-established roles in endothelial remodeling and immune cell trafficking. For example, CD44 is a key regulator of ECM remodeling, and CD44-deficient mice exhibit reduced collagenase activity and excessive fibrillar collagen deposition (Govindaraju *et al*, 2019). Endothelial CD44 also facilitates lymphocyte rolling and adhesion, linking matrix remodeling with immune cell recruitment (Nandi *et al*, 2000). Similarly, CD31 regulates both leukocyte transmigration and endothelial permeability through β-catenin–dependent junctional stabilization (Fernández-Martín *et al*, 2012). Increased expression of these molecules in the absence of galectin-1 positions this endogenous lectin as an important regulator of immune-endothelial interaction, important for ABMR.

Integrins also emerged as central nodes in both the *LGALS1* knockdown-dependent proteomic changes and the IFNγ–*LGALS1* interaction analysis, particularly ITGB1 and ITGB5. ITGB1 is a ubiquitous adhesion receptor that directly interacts with galectin-1 (Nam *et al*, 2017; Moiseeva *et al*, 1999), and galectin-1–integrin binding was shown to induce FAK phosphorylation and modulate cell migration (64, 65). Galectin-1 was previously linked to fibrosis and is important for repair of damaged tissues (Hermenean *et al*, 2022). Interestingly, in dermal fibrosis, galectin-1 colocalized with ITGB1, CD44, cadherins, and thrombospondin in fibroblasts, in association with immune infiltrates and collagen bundles (69), a phenotype reminiscent of the basement membrane thickening observed in chronic ABMR.

The interaction between IFNγ treatment and *LGALS1* knockdown further highlighted ITGB5 as a key integrator of inflammatory and adhesion signaling. Overexpression of ITGB5 was shown to activate Src and STAT3, leading to increased production of chemokines such as IL-8 and MCP1, which recruit neutrophils, monocytes, macrophages, NK cells, and lymphocytes (Leifheit-Nestler *et al*, 2010; Gschwandtner *et al*, 2019; Xiong *et al*, 2022). The identification of ITGB5 as an interaction-significant protein in our dataset suggests that galectin-1 could modulate IFNγ-driven immune recruitment through integrin-dependent signaling pathways. Additional focal adhesion and cytoskeletal adaptor proteins, including CD81, FSCN1, VE-cadherin, and ROBO4, were also altered, underscoring the breadth of galectin-1-dependent regulation of endothelial adhesion and ECM interaction. This alteration of ROBO4 is notable, as ROBO4 signaling has been found to limit Src-mediated VE-cadherin phosphorylation which leads to reduced cell-cell junctional adhesion (Koch *et al*, 2011). In sum, increase in cell-basement membrane adhesion molecules such as ITGB5, ITGB1, and CD44 due to *LGALS1* knockdown, but decrease in cell-cell adhesion molecules such as VE-cadherin and ROBO4 could suggest disruption of endothelial cell connections and may be responsible for weakening endothelial junctions and increased vascular permeability.

Phosphoproteomic analyses provided additional mechanistic resolution by implicating *LGALS1*- and IFNγ-dependent regulation of phosphorylation programs governing cytoskeletal organization, endocytosis, and caveolae formation, pathways that are central to endothelial mechano-transduction and barrier function. Altered CAVN1 phosphorylation and disrupted CAV1–CAVN1 co-localization suggest impaired caveolar assembly, a process tightly linked to endothelial cell polarization, migration, and mechano-transduction (Hill *et al*, 2012). These findings are notable, considering prior observations that CAV1 overexpression and increased caveolae formation are features of endothelial injury in chronic ABMR (Yamamoto *et al*, 2008; Nakada *et al*, 2016; Gambella *et al*, 2021), supporting the relevance of this pathway to disease-associated endothelial stress. Consistent with the role of galectin-1 in modulating caveolae-associated signaling, others have shown that stimulation with r-galectin-1 increased CAV1 phosphorylation (Lee *et al*, 2009). Moreover, ITGB1 engagement with ECM components such as fibronectin, an ECM protein that also interacts with galectin-1 (Ozeki *et al*, 1995), can induce phosphorylation of ITGB5, Src, FAK, and CAV1 (Park *et al*, 2011), highlighting the close interactions between adhesion- and caveolae-associated signaling networks. Taken together, these data raise the possibility that galectin-1-dependent perturbation of caveolar organization represents one mechanism linking altered adhesion signaling to endothelial barrier dysfunction, consistent with the increased permeability observed in our functional assays. These findings underscore the need for future studies to directly interrogate the functional consequences and temporal dynamics of these phosphorylation events.

Consistent with these molecular changes, functional studies using a perfusable microvascular model demonstrated that IFNγ disrupts GMEC barrier integrity and that *LGALS1* knockdown interacts with IFNγ to exacerbate this effect. In addition to changes in permeability, both IFNγ treatment and *LGALS1* silencing resulted in decreased secretion of GM-CSF, a cytokine with established roles in microvascular angiogenesis, endothelial repair, and barrier stabilization (Zhao *et al*, 2014; Thijssen, 2021). Endothelial cells typically exhibit low baseline GM-CSF expression (An *et al*, 2022), and in other cell types IFNγ suppressed GM-CSF production through STAT1-mediated signaling (Sharkey *et al*, 2018; Hu & Ivashkiv, 2009). The observation that *LGALS1* knockdown alone also reduced GM-CSF secretion suggests that intracellular galectin-1 positively regulates this angiogenic and barrier-supportive pathway, independently of IFNγ. Supporting this notion, the interplay between galectin-1 and GM-CSF has been implicated in angiogenesis (Blidner *et al*, 2025), and GM-CSF treatment accelerated wound healing and reduced vascular permeability (Yan *et al*, 2017). Moreover, *Lgals1* knockout models of tissue regeneration exhibited reduced expression of angiogenesis-related genes and increased expression of *Zfp36*, a negative regulator of GM-CSF mRNA stability, providing a plausible mechanistic link between *LGALS1* loss and diminished GM-CSF production (Potikha *et al*, 2016; Carballo *et al*, 2000). In contrast, IFNγ robustly increased IL-6 secretion, with no additional effect of *LGALS1* silencing, consistent with the proinflammatory actions of IFNγ and our transcript-level findings. These selective cytokine effects highlight the complexity of the functions of galectin-1 during inflammatory signaling. Rather than broadly suppressing inflammation, galectin-1 appears to preferentially support endothelial repair and angiogenic capacity while allowing classical IFNγ-driven inflammatory programs to proceed. Interestingly, the addition of extracellular r-galectin-1 produced unexpected effects on permeability and cytokine secretion compared with intracellular galectin-1 depletion, emphasizing that the addition of exogenous galectin-1 is not a simple replacement for the intracellular endogenous lectin. Extracellular galectin-1 may engage different receptors or signaling pathways than intracellular galectin-1, leading to divergent effects on endothelial behavior. For example, extracellular galectin-1 is known to bind to several receptors via N- and O-glycans, thus activating signaling pathways through distinct receptors (Sundblad *et al*, 2017; Rabinovich *et al*, 2002; Blidner *et al*, 2025). In fact, extracellular galectin-1 can activate downstream STAT1/3 signalling (Nambiar *et al*, 2019; Nam *et al*, 2017). This STAT1/3 activation could in turn result in the upregulation of proinflammatory cytokines (Khodarev *et al*, 2012; Metwally *et al*, 2020). The increase in pro-inflammatory cytokines was associated with diminished vascular permeability with exogenous galectin-1 in AngioPlate, suggesting that GMEC expression of adhesion and ECM proteins, such as CD44, CDH5, and ITGB5, are a critical determinant of vascular integrity in this context. Indeed, when intracellular galectin-1 was absent or IFNγ was added, exogenous galectin-1 resulted in increased vascular permeability, likely related to the decreased expression of VE-cadherin and ROBO4, and increased expression of adhesion and integrin proteins. While this may seem counterintuitive with the anti-inflammatory roles of galectin-1 in cancer (Mariño *et al*, 2023), others have shown that extracellular galectin-1 can act as a damage associated molecular pattern and ramp up immune system activation (Russo *et al*, 2021). While we selected this dose after careful consideration, one could propose that the effects in this system may differ at higher or lower doses. Thus, in the inflamed allograft microenvironment, where both intracellular regulation and extracellular galectin-1 exposure are likely altered, this duality will have important implications for disease progression and therapeutic targeting of galectin-1.

Hence, in this study we comprehensively established GMECs as the predominant cellular source of galectin-1 in the human kidney, including in the setting of ABMR, using orthogonal methods. Second, we leveraged both global and phosphoprotemics to decipher in-depth effects of galectin-1 manipulation in primary human GMECs. Third, the biological relevance of these findings is revealed by functional validation in a three-dimensional glomerular capillary model and extensive validation. AngioPlate platform enabled assessment of endothelial barrier integrity under defined perturbations, demonstrating the effects of *LGALS1* depletion, IFNγ stimulation and r-galectin-1 treatment, as well as their interaction. These functional data provide an important link between proteomic alterations and endothelial behavior relevant to glomerular endothelial integrity.

This study has some limitations. Other than the inclusion of data from ABMR biopsies to establish cellular localization, the mechanistic experiments were all performed *in vitro*. As such, the study does not directly demonstrate that the galectin-1-dependent proteomic and phosphoproteomic changes identified here occur in human ABMR tissue or are causal drivers of rejection pathology *in vivo*. In this regard, future studies are warranted to assess the effects of intracellular and extracellular galectin-1 on kidney allograft rejection *in vivo*.

In summary, our findings support a model in which galectin-1 acts as a central regulator of glomerular endothelial responses to inflammatory stress by integrating immune signaling with adhesion, cytoskeletal organization, and angiogenic support. Loss or dysregulation of galectin-1 shifts the endothelial phenotype toward impaired repair, increased permeability, and altered inflammatory cytokines, processes that are central to the pathogenesis of chronic ABMR. Distinct effects of intracellular depletion versus extracellular supplementation highlight the context-dependent nature of galectin-1 biology. Targeting galectin-1-dependent pathways or restoring endothelial repair mechanisms downstream of this lectin may therefore represent a novel strategy to mitigate chronic glomerular injury in kidney transplantation and in other glomerular diseases.

## Acknowledgements

Special thanks to Anthony Wu for technical assistance with MSstatsPTM software.

This work was supported by the Canadian Institutes of Health Research Project Grant (469957), Kidney Research Scientist Core Education and National Training (KRESCENT) grant (CIHR148204, KRES160004, KRES160005), University of Toronto CRAFT Project Award and the University Health Network (UHN) Foundation grants (579068260776, 579067450776, 579072310776) to AK. This work was also supported by the Frederick Banting and Charles Best Canada Graduate Scholarship-Master’s, the Institute of Medical Science Open Fellowship Award, and Queen Elizabeth II Graduate Scholarships in Science and Technology to AB. Finally, Barbara Mendelsohn was instrumental for generously funding this project, ensuring its completion. Computational biology analyses were supported in part by funding from Natural Sciences Research Council (NSERC RGPIN-2024-04314), CIHR (#519474), Canada Foundation for Innovation (CFI #225404, #30865), and Ontario Research Fund (RDI #34876, RE010-020) to IJ. The funders had no role in study design, data collection and analysis, decision to publish, or preparation of the manuscript.

## Data availability

The data from this manuscript is available at ProteomeXchange with the identifiers PXD074919 (Proteome) and PXD074969 (Phosphoproteome). All other data are included in the manuscript or the Supplement.

## Supplemental data

This article contains supplemental data.

## Abbreviations

ABMR: Antibody-mediated rejection
GMEC: glomerular microvascular endothelial cells
IFNγ: Interferon gamma
r-galectin-1: recombinant galectin-1
*LGALS1*: galectin-1 gene
siRNA: small interfering RNA
NT_V: non-targeting control and vehicle
NT_IFNγ: non-targeting control and IFNγ Treatment
siRNA_V: *LGALS1* siRNA and vehicle
siRNA_IFNγ: *LGALS1* siRNA and IFNγ Treatment
GO-BP: Gene Ontology – Biological Processes
GO-MF: Gene Ontology – Molecular Function
GO-CC: Gene Ontology – Cellular Component
IID: Integrated Interactions Database
pathDIP: pathway Data Integration Portal
NAViGaTOR: Network Analysis, Visualization, & Graphing TORonto.

