## Supplemental Figures for "Galectin-1 Modulates Cell Adhesions, Caveolae, and Vascular Permeability in Kidney Endothelial Cells – Insights from Proteomics, Phosphoproteomics, and Functional Studies"

#### SUPPLEMENTAL FIGURE LEGENDS

**Supplemental Figure S1. Protein expression of Phospho-STAT1 and total STAT1 following IFNγ treatment of glomerular microvascular endothelial cells**

Western Blotting of Phospho-STAT1 (P-STAT1) (Y701) **(A)** and total-STAT1 **(B)** after 50U/mL of IFNγ or vehicle treatment for 30 minutes, following *LGALS1* siRNA knockdown or non-targeting control treatment. NT_V, non-targeting control and vehicle; NT_IFNγ, non-targeting control and IFNγ Treatment; siRNA_V, *LGALS1* siRNA and vehicle; siRNA_IFNγ, *LGALS1* siRNA and IFNγ treatment.

**Supplemental Figure S2. Protein expression of HLA Class II-DR following IFNγ treatment of glomerular microvascular endothelial cells**

Western blotting of HLA Class II-DR and vinculin housekeeping after 50U/mL of IFNγ or vehicle treatment for 24 hours. V, vehicle; IFNγ, interferon-Gamma treatment.

**Supplemental Figure S3. Histograms of total protein distribution in glomerular microvascular endothelial cell lysates after imputation**

Histograms depicting the distribution of the measured (blue) and imputed (red) protein intensity values in the total proteome, for each sample.

**Supplemental Figure S4. Imaging Mass Cytometry of biopsies from kidneys with ABMR**

Imaging mass cytometry images of 3 regions of interest from 3 biopsies with ABMR, stained for galectin-1 (red) and co-stained with **(A)** CD31 (green) or **(B)** CD45 (green), and nuclei (blue).

**Supplemental Figure S5. Imaging Mass Cytometry of living donor kidney biopsy**

Imaging mass cytometry images from 1 biopsy from a healthy living donor control, stained for galectin-1 (red) and co-stained with **(A)** CD31 (green) or **(B)** CD45 (green), and nuclei (blue).

**Supplemental Figure S6. Characterization of GMEC phenotypic gene expression and gene expression associated with GMEC activation following *LGALS1* silencing**

**A)** Gene expression of *LGALS1* at 48, 72, and 96 hours post *LGALS1* silencing compared to non-targeting control. **B)** Galectin-1 protein concentration in GMEC supernatant, 72 hours post *LGALS1* knockdown. **C)** Gene expression of *PECAM1*, *CDH5*, and *VWF* at 48, 72, and 96 hours post *LGALS1* knockdown. **D)** Gene expression of *ACTA2, IL6*, and *ICAM1* at 48, 72, and 96 hours post *LGALS1* knockdown. Values normalized to *ACTB* housekeeping gene and relative to NT control at each timepoint. Data presented as mean ±SEM, n = 3-6 per group. Mann-Whitney U Test, $^*P<0.05$, $^{***}P<0.001$. NT, non-targeting control; LGALS1 siRNA, siRNA targeting *LGALS1* gene.

**Supplemental Figure S7. Pathways enriched among proteins increased and decreased with IFNγ treatment**

Bubble plots of pathways significantly enriched among proteins differentially expressed with IFNγ treatment compared to vehicle control. **A)** Plot of the top 10 enriched pathways among proteins increased with IFNγ treatment compared to vehicle control (FDR: BH<0.01). **B)** Plot of the top 10 enriched pathways among proteins decreased with IFNγ treatment compared to vehicle control

(FDR: BH<0.01). The size of the circle indicates the number of proteins in that pathway and the colour indicates significance.

**Supplemental Figure S8. Heatmap of significantly differentially expressed proteins in *LGALS1* siRNA compared to non-targeting control**

Full unsupervised hierarchical heatmap of 236 proteins that were significantly differentially expressed with *LGALS1* siRNA compared to NT control. Red, green, blue, and yellow boxes denote cluster 1, 2, 3, and 4, respectively. 2-way ANOVA with Tukey's post-test, $p<0.05$. NT control, non-targeting control.

**Supplemental Figure S9. Heatmap of proteins significant for the interaction of *LGALS1* siRNA and IFNγ treatment**

Full unsupervised hierarchical heatmap of 267 proteins that were significantly differentially expressed for the interaction of *LGALS1* siRNA and IFNγ treatment. 2-way ANOVA with Tukey's post-test, $p<0.05$.

**Supplemental Figure S10. *LGALS1* silencing and IFNγ treatment alter ITGB5 protein expression in GMECs**

Western Blotting of β5 integrin (ITGB5) **(A)** and GAPDH as housekeeping control **(B)** GMECs treated with *LGALS1* siRNA, IFNγ, or both. V, vehicle; IFNγ, Interferon-gamma treatment; NT_V, non-targeting control and vehicle; NT_IFNγ, non-targeting control and IFNγ Treatment; siRNA_V, *LGALS1* siRNA and vehicle; siRNA_IFNγ, *LGALS1* siRNA and IFNγ treatment.

**Supplemental Figure S11. *LGALS1* silencing and IFNγ treatment cause loss of CAVN1/CAV1 co-localization in GMECs**

Immunostaining of CAVN1 and CAV1 when GMECs were treated with *LGALS1* siRNA, IFNγ, or both. Blue colour is DAPI, green colour stains CAV1, red colour stains CAVN1. Red arrows denote yellow co-staining of CAV1 and CAVN1 in the NT_V condition. NT_V, non-targeting control and vehicle; NT_IFNγ, non-targeting control and IFNγ treatment; siRNA_V, *LGALS1* siRNA and vehicle; siRNA_IFNγ, *LGALS1* siRNA and IFNγ treatment.

**Supplemental Figure S12. *LGALS1* silencing, recombinant galectin-1, and IFNγ treatment alter GMEC vascular permeability to 65kDa Dextran and cytokine secretion**

**A)** GMEC vessel barrier integrity was assessed by diffusion of 65kDa dextran in μg/hour after treatment with IFNγ, *LGALS1* siRNA, both, or NT and vehicle control. Data presented as mean ±SEM, n=15-16 per group, Aligned Rank Sum Test, ***P<0.001 ****P<0.0001. **B)** IFNγ, IL-8, and MCP-1 cytokine concentrations in flowthrough from wells treated with *LGALS1* siRNA, IFNγ, both, or NT and vehicle control. Data presented as mean ±SEM, n=5-6 per group, Aligned Rank Sum Test, *P<0.05, ***P<0.001, ****P<0.0001. **C)** GMEC vessel barrier integrity was assessed by diffusion of 65kDa dextran in μg/hour after treatment with *LGALS1* siRNA, IFNγ, both, or NT and vehicle control, followed by the addition of r-galectin-1 or vehicle. Data are presented as mean ±SEM, n=7-8 per group, Aligned Rank Sum Test, *P<0.05, ***P<0.001. **D)** IFNγ, IL8, and MCP-1 cytokine concentration in flowthrough from wells treated with *LGALS1* siRNA, IFNγ, both, or NT and vehicle control, followed by the addition of r-galectin-1 or vehicle control. Data are presented as mean ±SEM, n=5-6 per group, Aligned Rank Sum Test, *P<0.05, ***P<0.001, ****P<0.0001. NT, non-targeting control and vehicle; NT_r-galectin-1, non-targeting control and vehicle and recombinant galectin-1; NT_IFNγ, non-targeting control and IFNγ treatment; NT_IFNγ_r-galectin-1, non-targeting control and IFNγ treatment and recombinant galectin-1; siRNA, *LGALS1* siRNA and vehicle; siRNA_r-galectin-1, *LGALS1* siRNA and vehicle and recombinant galectin-1; siRNA_IFNγ, *LGALS1* siRNA and IFNγ treatment; siRNA_IFNγ_r-galectin-1, *LGALS1* siRNA and IFNγ treatment and recombinant galectin-1.

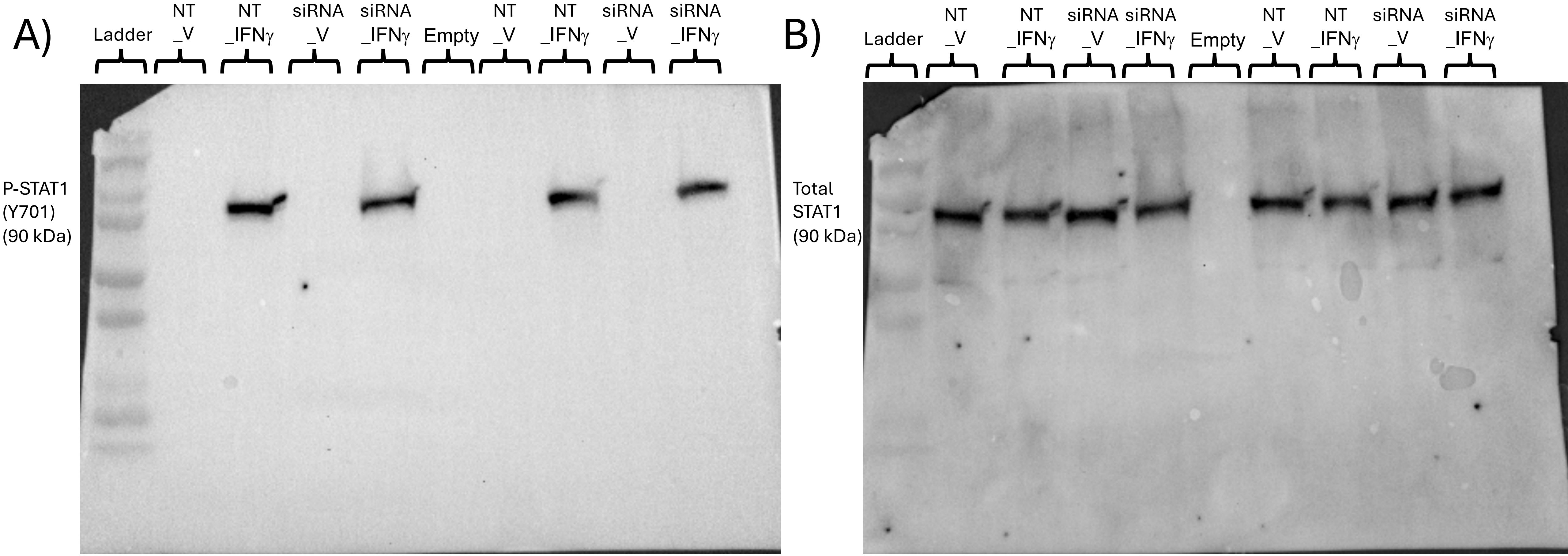
A)
Ladder
NT _V
NT _IFNγ
siRNA _V
siRNA _IFNγ
Empty
NT _V
NT _IFNγ
siRNA _V
siRNA _IFNγ
P-STAT1 (Y701) (90 kDa)
B)
Ladder
NT _V
NT _IFNγ
siRNA _V
siRNA _IFNγ
Empty
NT _V
NT _IFNγ
siRNA _V
siRNA _IFNγ
Total STAT1 (90 kDa)

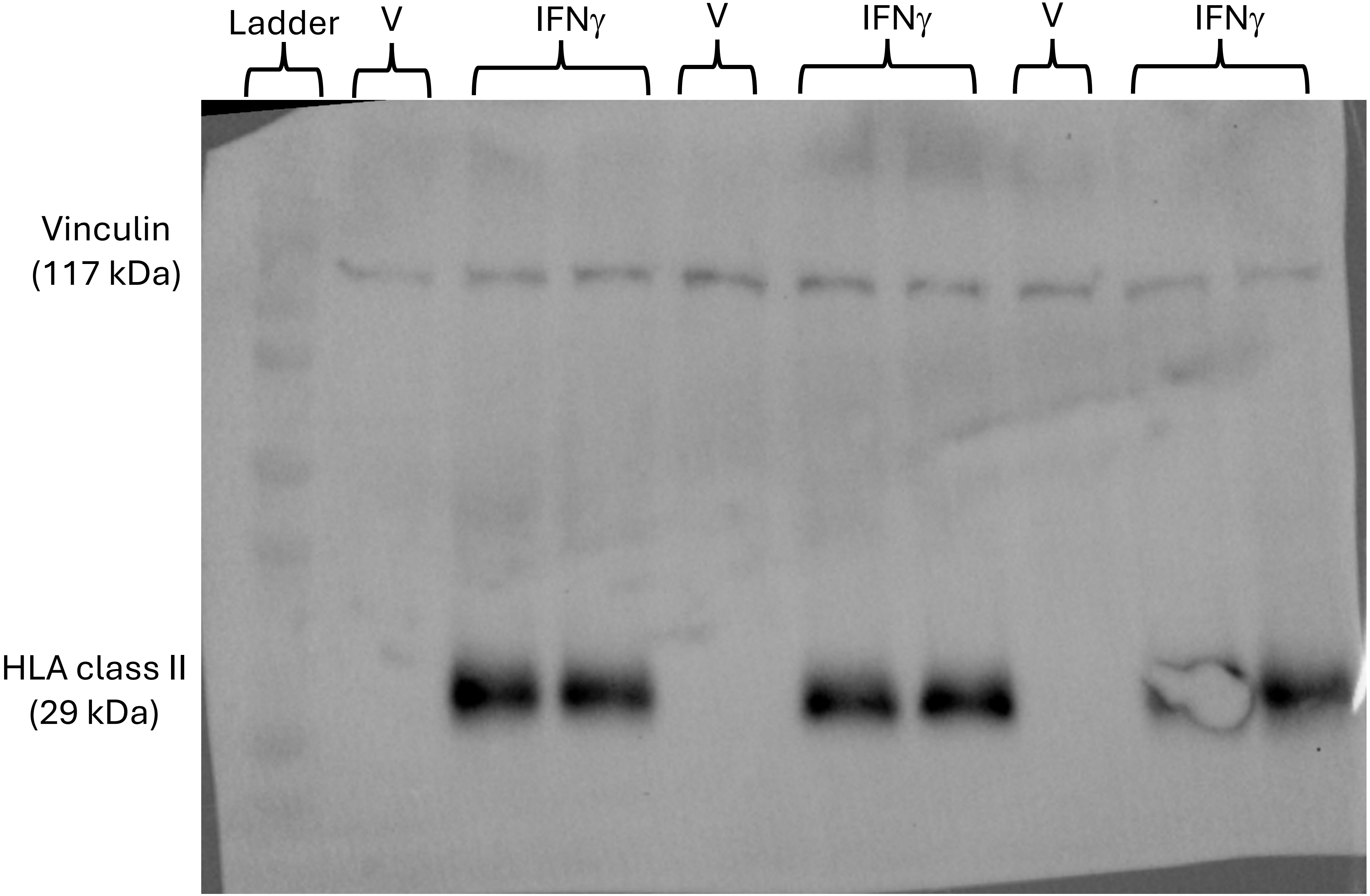

Ladder
V
IFNγ
V
IFNγ
V
IFNγ
Vinculin
(117 kDa)
HLA class II
(29 kDa)

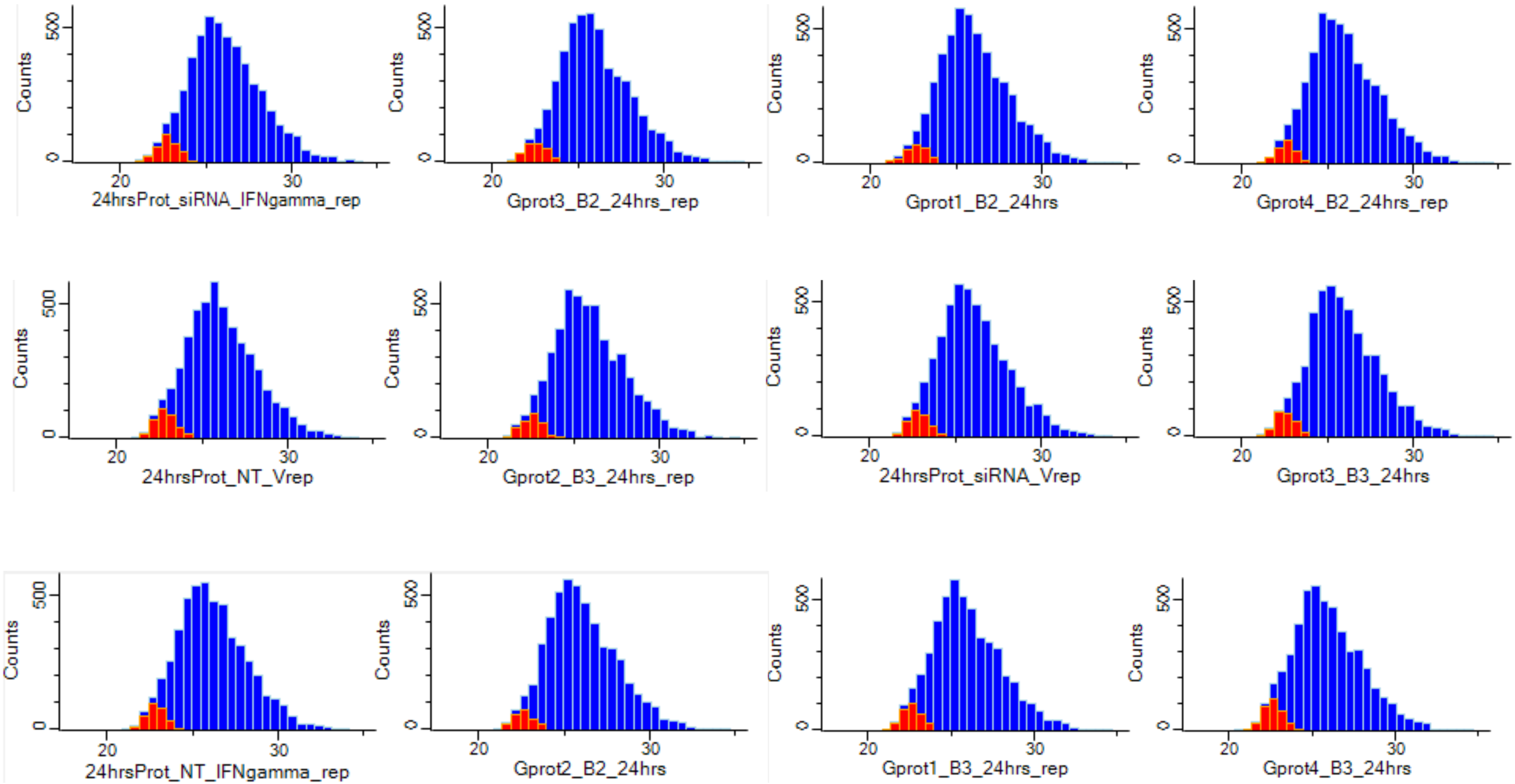
Counts
500
0
20
30
24hrsProt_siRNA_IFNgamma_rep
Gprot3_B2_24hrs_rep
Gprot1_B2_24hrs
Gprot4_B2_24hrs_rep
24hrsProt_NT_Vrep
Gprot2_B3_24hrs_rep
24hrsProt_siRNA_Vrep
Gprot3_B3_24hrs
24hrsProt_NT_IFNgamma_rep
Gprot2_B2_24hrs
Gprot1_B3_24hrs_rep
Gprot4_B3_24hrs

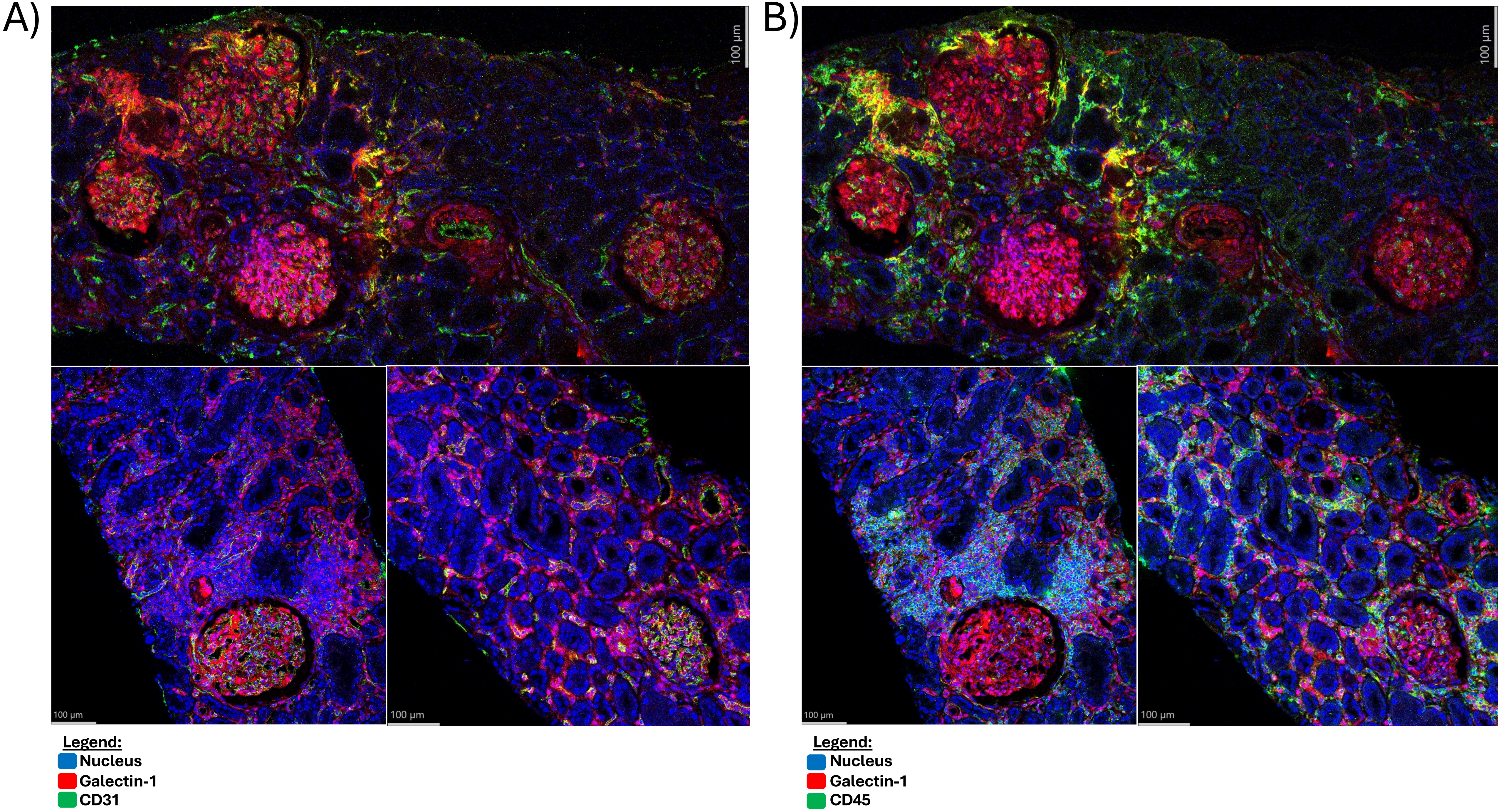
A)
B)
100 µm
100 µm
100 µm
100 µm
100 µm
100 µm
Legend:
Nucleus
Galectin-1
CD31
Legend:
Nucleus
Galectin-1
CD45

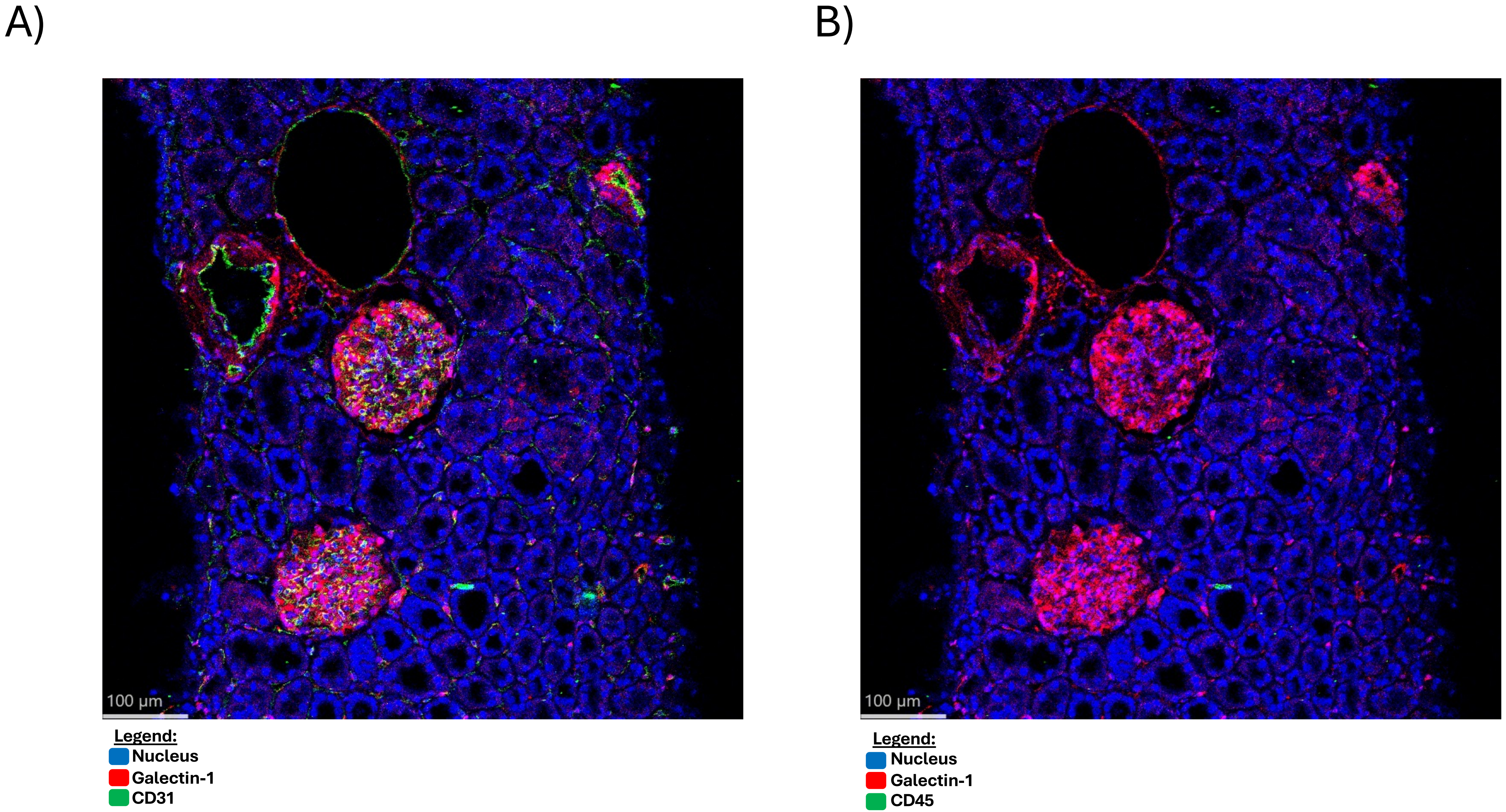
A)
100 µm
Legend:
Nucleus
Galectin-1
CD31
B)
100 µm
Legend:
Nucleus
Galectin-1
CD45

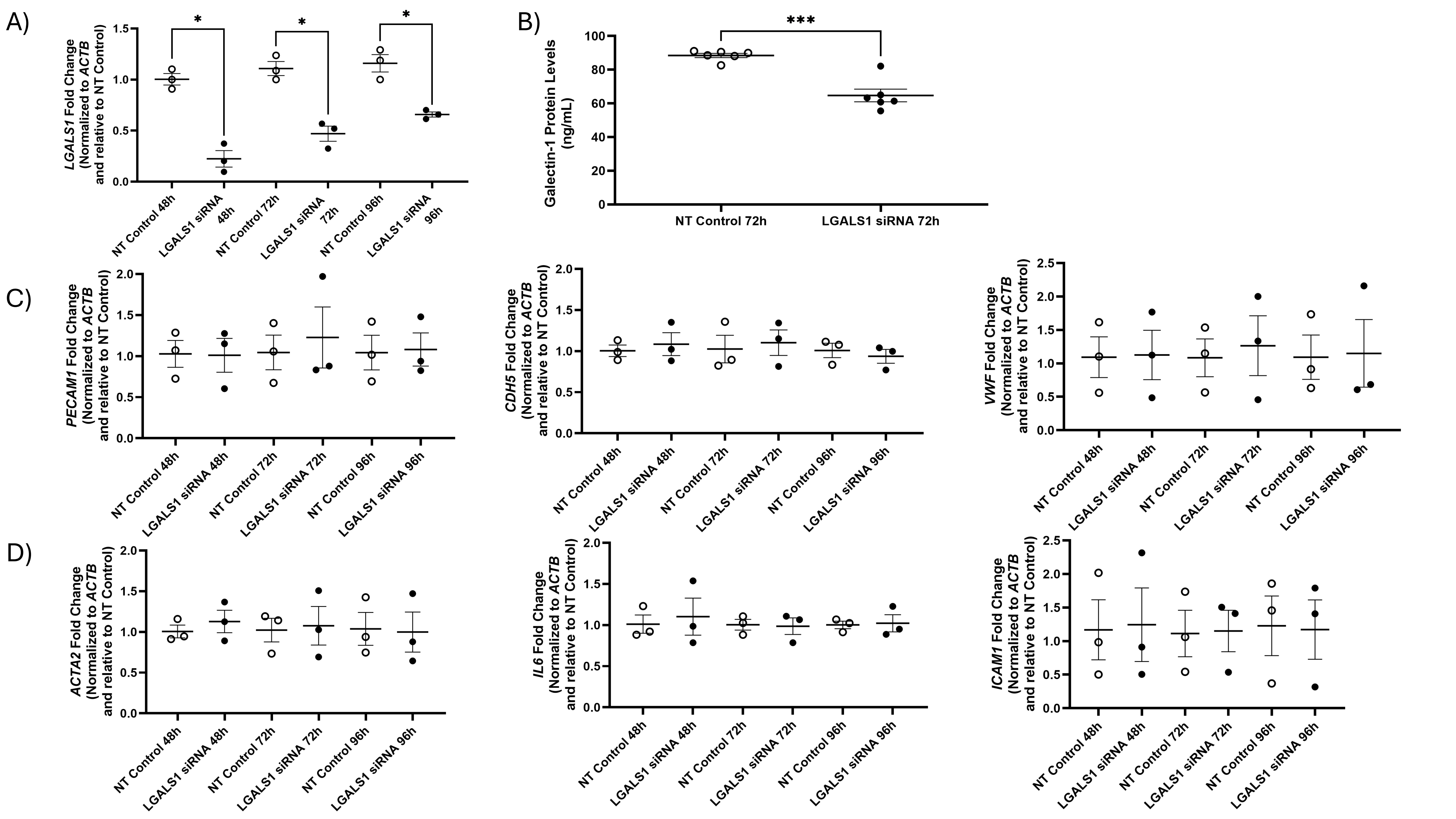
A)
LGALS1 Fold Change (Normalized to ACTB and relative to NT Control)
NT Control 48h
LGALS1 siRNA 48h
NT Control 72h
LGALS1 siRNA 72h
NT Control 96h
LGALS1 siRNA 96h
B)
Galectin-1 Protein Levels (ng/mL)
***
NT Control 72h
LGALS1 siRNA 72h
C)
PECAM1 Fold Change (Normalized to ACTB and relative to NT Control)
CDH5 Fold Change (Normalized to ACTB and relative to NT Control)
VWF Fold Change (Normalized to ACTB and relative to NT Control)
D)
ACTA2 Fold Change (Normalized to ACTB and relative to NT Control)
IL6 Fold Change (Normalized to ACTB and relative to NT Control)
ICAM1 Fold Change (Normalized to ACTB and relative to NT Control)

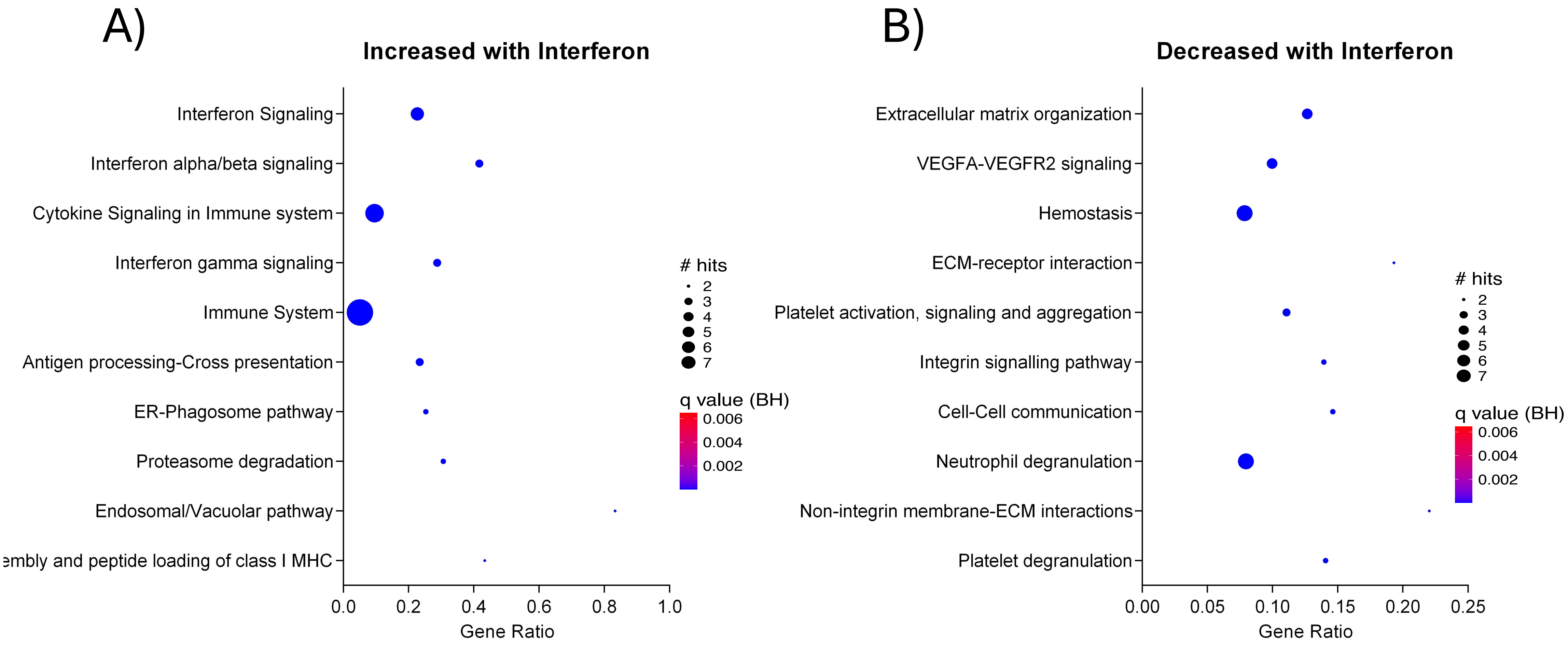
A)
Increased with Interferon
Interferon Signaling
Interferon alpha/beta signaling
Cytokine Signaling in Immune system
Interferon gamma signaling
Immune System
Antigen processing-Cross presentation
ER-Phagosome pathway
Proteasome degradation
Endosomal/Vacuolar pathway
embly and peptide loading of class I MHC
0.0
0.2
0.4
0.6
0.8
1.0
Gene Ratio
### hits
2
3
4
5
6
7
q value (BH)
0.006
0.004
0.002
B)
Decreased with Interferon
Extracellular matrix organization
VEGFA-VEGFR2 signaling
Hemostasis
ECM-receptor interaction
Platelet activation, signaling and aggregation
Integrin signalling pathway
Cell-Cell communication
Neutrophil degranulation
Non-integrin membrane-ECM interactions
Platelet degranulation
0.00
0.05
0.10
0.15
0.20
0.25
Gene Ratio
### hits
2
3
4
5
6
7
q value (BH)
0.006
0.004
0.002

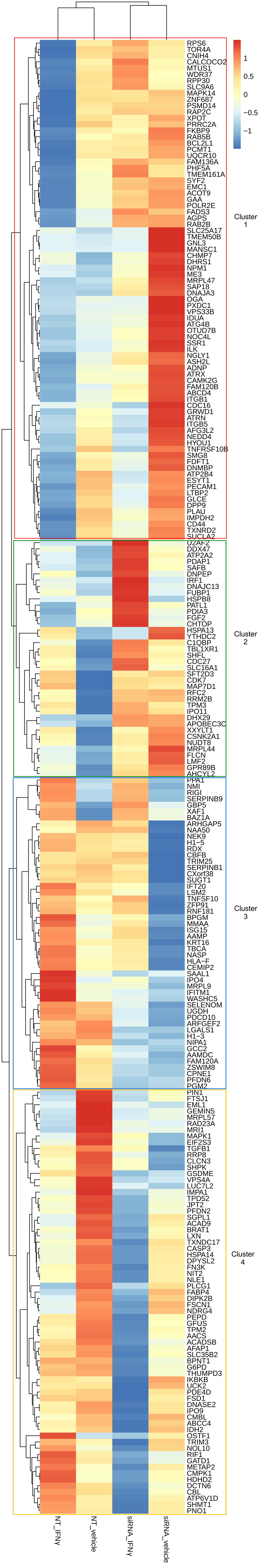
1
0.5
0
−0.5
−1
Cluster 1
Cluster 2
Cluster 3
Cluster 4
RPS6
TOR4A
CNIH4
CALCOCO2
MTUS1
WDR37
RPP30
SLC9A6
MAPK14
ZNF687
PSMD14
RAP2C
XPOT
PRRC2A
FKBP9
RAB5B
BCL2L1
PCMT1
UQCR10
FAM136A
PHF5A
TMEM161A
SYF2
EMC1
ACOT9
GAA
POLR2E
FADS3
AGPS
RAB2B
SLC25A17
TMEM50B
GNL3
MANSC1
CHMP7
DHRS1
NPM1
ME3
MRPL47
SAP18
DNAJA3
OGA
PXDC1
VPS33B
IDUA
ATG4B
OTUD7B
NOC4L
SSR1
ILK
NGLY1
ASH2L
ADNP
ATRX
CAMK2G
FAM120B
ABCD4
ITGB1
CDC16
GRWD1
ATRN
ITGB5
AFG3L2
NEDD4
HYOU1
TNFRSF10B
SMG8
FDFT1
DNMBP
ATP2B4
ESYT1
PECAM1
LTBP2
GLCE
DPP9
PLAU
IMPDH2
CD44
TXNRD2
SUCLA2
U2AF2
DDX47
ATP2A2
PDAP1
SAFB
DNPEP
IRF1
DNAJC13
FUBP1
HSPB8
PATL1
PDIA3
FGF2
CHTOP
HSPA13
YTHDC2
C1QBP
TBL1XR1
SHFL
CDC27
SLC16A1
SFT2D3
CDK7
MAP7D1
RFC2
RRM2B
TPM3
IPO11
DHX29
APOBEC3C
XXYLT1
CSNK2A1
NUDT8
MRPL44
FLCN
LMF2
GPR89B
AHCYL2
PPA1
NMI
RIGI
SERPINB9
GBP5
XAF1
BAZ1A
ARHGAP5
NAA50
NEK9
H1−5
RDX
CBFB
TRIM25
SERPINB1
CXorf38
SUGT1
IFT20
LSM2
TNFSF10
ZFP91
RNF181
BPGM
MMAA
ISG15
AAMP
KRT16
TBCA
NASP
HLA−F
CEMIP2
SAAL1
IPO4
MRPL9
IFITM1
WASHC5
SELENOM
UGDH
PDCD10
ARFGEF2
LGALS1
H1−3
NIPA1
GCC2
AAMDC
FAM120A
ZSWIM8
CPNE1
PFDN6
PGM2
PIN1
FTSJ1
EML1
GEMIN5
MRPL57
RAD23A
MRI1
MAPK1
EIF2S3
TGFB1
RRP8
CLCN3
SHPK
GSDME
VPS4A
LUC7L2
IMPA1
TPD52
JPT2
PFDN2
SGPL1
ACAD9
BRAT1
LXN
TXNDC17
CASP3
HSPA14
DPYSL2
FN3K
NIT2
NLE1
PLCG1
FABP4
DIPK2B
FSCN1
NDRG4
PEPD
GFUS
TPM2
AACS
ACADSB
AFAP1
SLC35B2
BPNT1
G6PD
THUMPD3
IKBKB
UCK2
PDE4D
FSD1
DNASE2
IPO9
CMBL
ABCC4
IDH2
OSTF1
TRIM3
NOL10
RIF1
GATD1
METAP2
CMPK1
HDHD2
DCTN6
CBL
ATP6V1D
SHMT1
PNO1
NT_IFNγ
NT_vehicle
siRNA_IFNγ
siRNA_vehicle

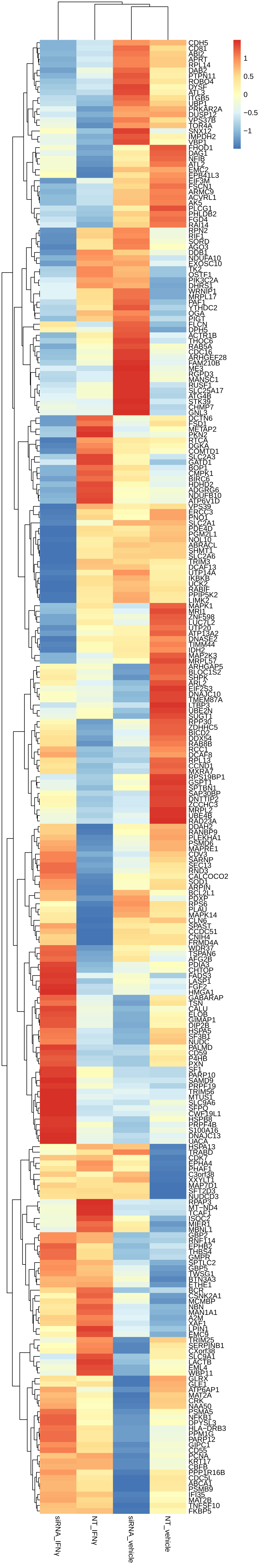
1
0.5
0
−0.5
−1
CDH5
CD81
ABI2
APRT
RPL14
DAB2
PTPN11
ROBO4
DYSF
ATL3
ITGB5
UBP1
PRKAR2A
DUSP12
VPS37B
TOR4A
SNX12
IMPDH2
VBP1
FHOD1
DAG1
NFIB
ATL2
EMC2
EPB41L3
EIF3M
FSCN1
ARMC9
ACVRL1
AK5
PLCG1
PHLDB2
FGD4
RAI14
RPN2
RIF1
SORD
ACO2
DDB1
NDUFA10
EXOSC10
TK2
OSTF1
PIK3C2A
DHRS1
WRNIP1
MRPL17
PAF1
YTHDC2
OGA
PIGT
FLCN
DPH5
ACTR1B
THOC6
RAB5A
CDC16
ARHGEF28
FAM210B
ME3
RGPD3
MANSC1
RUSF1
SLC25A17
ATG4B
STK39
CHMP7
GNL3
DCTN6
FSD1
METAP2
PKN2
RTCA
DGKA
COMTD1
SLC2A3
GATD1
BOP1
CMPK1
BIRC6
HDHD2
ADGRG6
NDUFB10
ATP6V1D
VPS39
ERCC3
PNO1
SLC2A1
PDE4D
PGM2L1
NOL10
ABRACL
SHMT1
SLC2A6
TRIM3
DCAF13
UTP14A
IKBKB
UCK2
RABIF
PPIP5K2
LIMK2
MAPK1
MRI1
ZNF598
LUC7L2
UTP20
ATP13A2
DNASE2
TIMM44
IDH2
MAP2K3
MRPL57
ARHGAP5
BLOC1S2
SHPK
ARL2
EIF2S3
DNAJC10
TMEM87A
LTBP3
UBE2N
SUGT1
RPP30
ZDHHC5
BICD2
DDX54
RAB8B
RCC1
DCAF8
RPL13
CCND1
MXRA7
RPS19BP1
GSPT1
SPTBN1
SAP30BP
DNTTIP2
ZCCHC3
MRPL2
UBE4B
RAD23A
DDAH2
RANBP9
PLEKHA1
PSMD6
MAPRE1
CDV3
SARNP
SEC13
RND3
CALCOCO2
SOD1
ARPIN
BCL2L1
PDXP
RPS6
PLAU
MAPK14
CLN6
SPAST
CCDC51
CNIH4
FRMD4A
WDR37
TSPAN6
AFG2B
PDIA3
CHTOP
FADS3
LASP1
FGF2
HMGA1
GABARAP
TSN
CALU
ELOB
GIMAP1
DIP2B
HSPA5
SF3B1
NUDC
PALMD
CD59
P4HB
PXN
SF1
PARP10
SAMD9
PRPF19
TRIM56
MTUS1
SLC9A6
SFPQ
CWF19L1
HSPB8
PRPF4B
S100A16
DNAJC13
UACA
HSPA13
TRABD
CDK7
EPHA4
PHAF1
C3orf38
XXYLT1
MAP7D1
SFT2D3
NUDCD3
RPAP3
MT-ND4
TCAF1
ISOC2
MIER1
MBNL1
GBP2
RNF114
EPHB2
THBS4
GMPR
SPTLC2
GBP5
TWSG1
BTN3A3
ETHE1
BCR
CSNK2A1
MCMBP
NBN
MAN1A1
A2M
XAF1
LPIN1
EMC9
TRIM25
SERPINB1
CXorf38
SLC9A1
LACTB
EML4
WBP11
GLRX
GLE1
ATP6AP1
MAT2A
CRK
NAA50
PSMA5
NFKB1
DPYSL3
HLA-DRB3
PPM1G
PARP12
GIPC1
CD55
PCNA
KRT17
CBFB
PPP1R16B
CDC5L
ABCA1
PSMB9
IFI35
MAT2B
TNFSF10
FKBP5
siRNA_IFNγ
NT_IFNγ
siRNA_vehicle
NT_vehicle

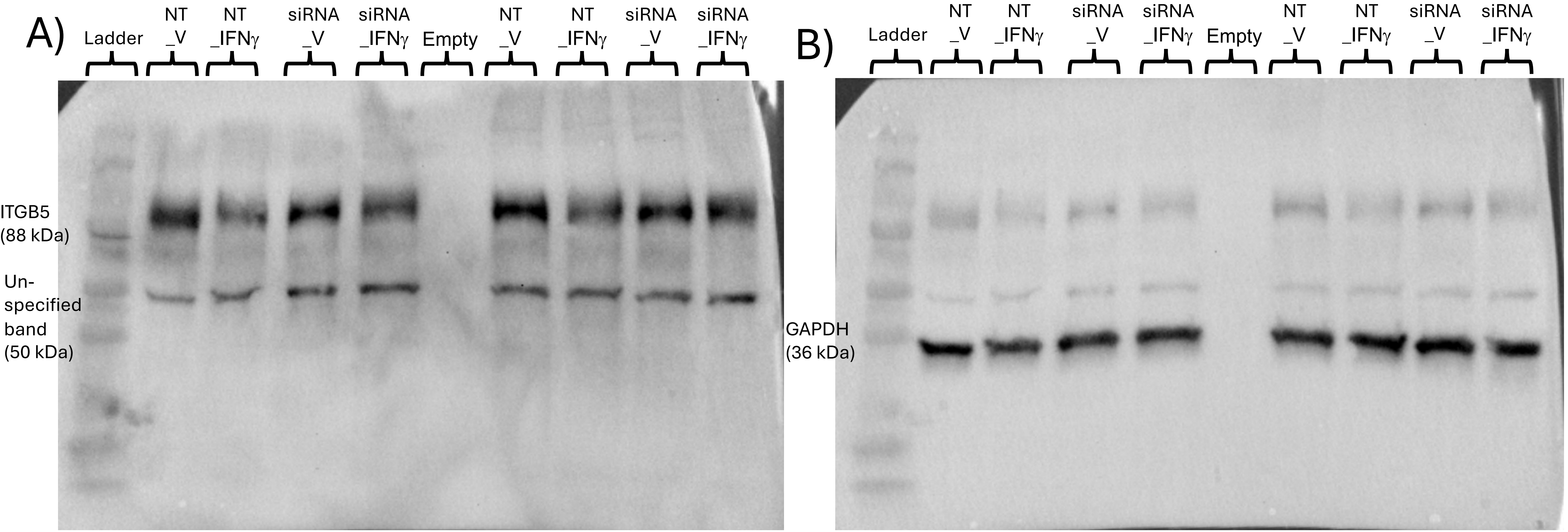
A)
Ladder
NT _V
NT _IFNγ
siRNA _V
siRNA _IFNγ
Empty
NT _V
NT _IFNγ
siRNA _V
siRNA _IFNγ
ITGB5 (88 kDa)
Un-specified band (50 kDa)
B)
Ladder
NT _V
NT _IFNγ
siRNA _V
siRNA _IFNγ
Empty
NT _V
NT _IFNγ
siRNA _V
siRNA _IFNγ
GAPDH (36 kDa)

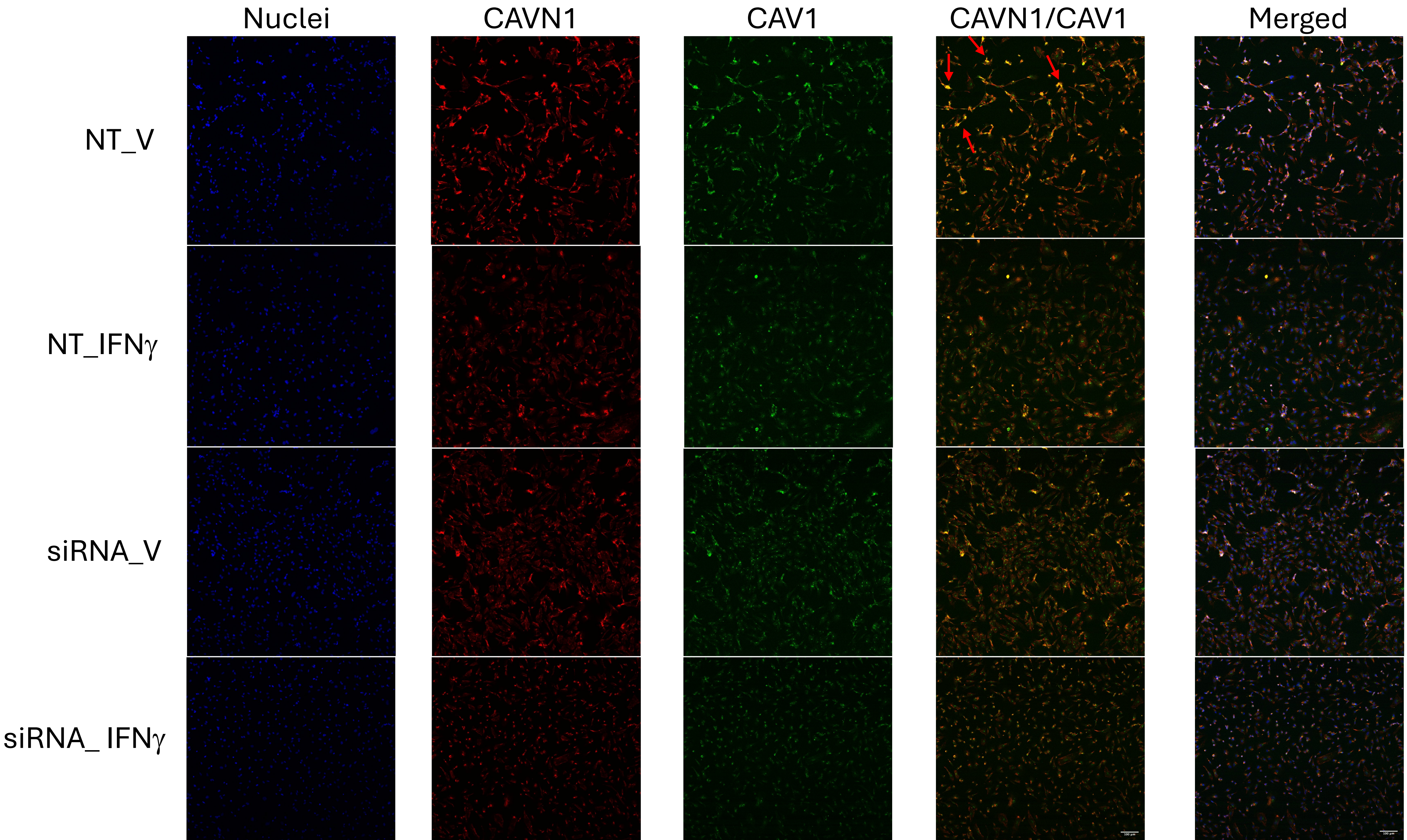
Nuclei
CAVN1
CAV1
CAVN1/CAV1
Merged
NT_V
NT_IFNγ
siRNA_V
siRNA_ IFNγ

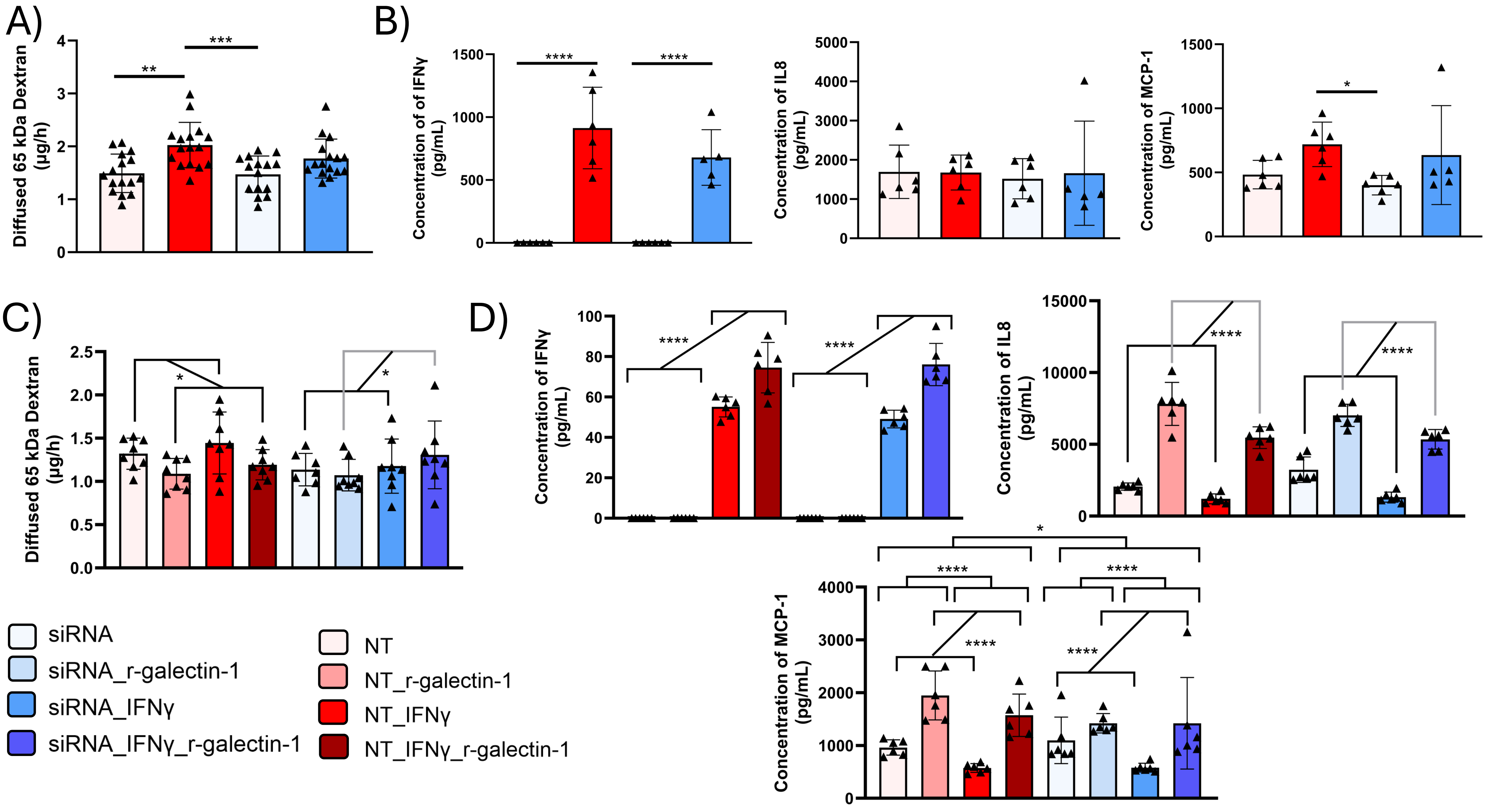
A)
B)
C)
D)
Diffused 65 kDa Dextran (µg/h)
Concentration of of IFNγ (pg/mL)
Concentration of IL8 (pg/mL)
Concentration of MCP-1 (pg/mL)
Concentration of IFNγ (pg/mL)
siRNA
siRNA_r-galectin-1
siRNA_IFNγ
siRNA_IFNγ_r-galectin-1
NT
NT_r-galectin-1
NT_IFNγ
NT_IFNγ_r-galectin-1
